# The LexA-RecA* structure reveals a lock-and-key mechanism for SOS activation

**DOI:** 10.1101/2023.10.30.564768

**Authors:** Michael B. Cory, Allen Li, Christina M. Hurley, Peter J. Carman, Ruth A. Pumroy, Zachary M. Hostetler, Yarra Venkatesh, Kushol Gupta, E. James Petersson, Rahul M. Kohli

## Abstract

The bacterial SOS response plays a key role in adaptation to DNA damage, including that caused by antibiotics. SOS induction begins when activated RecA*, an oligomeric nucleoprotein filament formed on single-stranded DNA, binds to and stimulates autoproteolysis of the repressor LexA. Here, we present the structure of the complete SOS signal complex, constituting full-length LexA bound to RecA*. We uncover an extensive interface unexpectedly including the LexA DNA-binding domain, providing a new molecular rationale for ordered SOS response gene induction. Furthermore, we find that the interface involves three RecA monomers, with a single residue in the central monomer acting as a molecular key, inserting into an allosteric binding pocket to induce LexA cleavage. Given the pro-mutagenic nature of SOS activation, our structural and mechanistic insights provide a foundation for developing new therapeutics to slow the evolution of antibiotic resistance.

## INTRODUCTION

Infections caused by antibiotic resistant bacterial pathogens represent an increasingly dire threat. This clinical problem is due, in part, to the remarkable ability of bacteria to sense, respond, and adapt to environmental stresses, including those posed by antibiotics. Bacterial adaptation to antimicrobial stress can occur in multiple ways, including acquisition of resistance-conferring mutations, transfer of resistance genes, or induction of protective states such as biofilms^1–3^. Many of these evasion mechanisms are part of a generalized DNA damage-inducible genetic network, known as the SOS response^4–6^. Indeed, genetic studies have confirmed that inactivation of the SOS response can render bacteria hypersensitive to DNA damaging antibiotics^7–10^, while simultaneously slowing the acquisition of resistance^9,11–13^. Together, these studies support the proposal that SOS inhibition could strengthen our antibiotic arsenal in two ways: synergy with DNA damaging antibiotics by increasing potency and suppression of acquired resistance in bacteria that survive initial treatment.

The SOS response is under the control of two well-conserved regulatory proteins, LexA and RecA. LexA is a homodimer with each monomer containing an N-terminal DNA-binding domain (NTD) and a C-terminal serine protease domain (CTD). Dimeric LexA recognizes and represses SOS response gene promoters, which number more than 50 in *E. coli*^14–17^. The initiating molecular signal for SOS activation is the accumulation of single-stranded DNA (ssDNA), typically due to replication stalling at sites of DNA damage^18–22^. This ssDNA acts as a template for the ATP-dependent polymerization of RecA into extended, helical nucleoprotein filaments, referred to as RecA*. While RecA* acts in pathways of homologous recombination, it also serves a key secondary role as a ‘co-protease’ for LexA^4,18^. When LexA binds to RecA*, a conformational change is induced in the LexA CTD that brings the catalytic serine-lysine dyad into proximity with an internal protein cleavage loop containing a scissile bond^14,23–29^. Autoproteolysis severs the DNA-binding NTD domain from the CTD dimerization interface, destabilizing LexA and thereby driving de-repression of SOS response genes^16,17,30^ (**Fig. 1A**).

**Figure 1.**
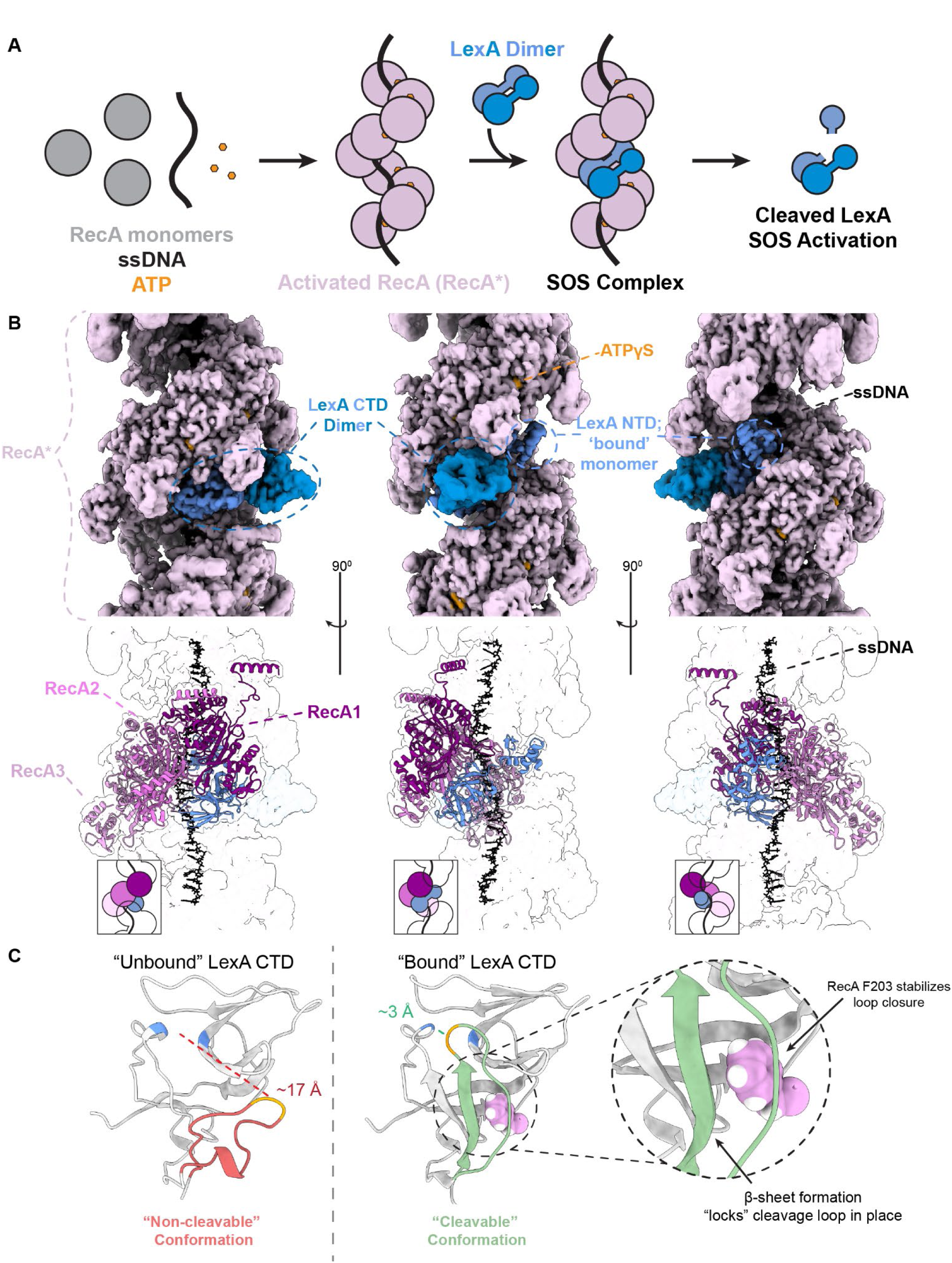
Structure of the SOS signal complex. **A)** Cartoon schematic of SOS activation, highlighting RecA filamentation, SOS complex formation, and LexA proteolysis. SOS activation is associated with the ordered induction of DNA damage response genes, including those assocaited with acquired antibiotic resistance. **B)** At top, density of SOS complex shown as a solid surface. Major structural features of the complex are highlighted and labeled. At bottom, map density is shown as a contour, with ribbons displaying a timer of RecA units (pink, magenta, purple) engaged with the bound LexA monomer (blue), with ssDNA (black) as the filament axis. Insets show a cartoon depiction of the oriented filament for each view. **C)** Comparison of the “unbound” and “bound” LexA CTDs within the dimer. The cleavage loop is colored in either red or green, with the target scissile bond highlighted in orange and the catalytic serine protease active site residues highlighted in blue. Distances from the target bond to the hydroxyl of the catalytic serine are given. A single RecA residue, F203 from the RecA2 unit, is shown in a space-filling model (pink) associated with the bound LexA monomer.

Formation of the ‘SOS signal complex’ – the ternary complex between RecA, ssDNA and dimeric full-length LexA – thus serves as the key regulatory step governing the SOS response^31^. A remarkable aspect of the SOS response is that gene induction is orchestrated in a chronological fashion, despite the fact that LexA serves as the only negative regulator governing expression^32–34^. While early response elements include high-fidelity DNA repair proteins, such as nucleotide excision repair enzymes^4,34,35^, late response elements include DNA damage tolerance genes, such as pro-mutagenic translesion synthesis DNA polymerases^34–37^. This dual role provides a rationale for the observations that SOS inactivation can result in both hypersensitization to DNA damaging antibiotics and decreased acquisition of antibiotic resistance^9^. Beyond the necessity of SOS for repairing and tolerating DNA damage, the SOS response has been shown to govern other adaptive mechanisms such as mobile resistance gene expression in *V. cholerae*, CRISPR-induced mutagenesis in *S. epidermidis*, and biofilm formation in P*. aeruginosa*^6,38–40^.

Despite decades of study, the absence of insight into the structural underpinnings of SOS complex formation has been one limitation to the rational design of small-molecule SOS antagonists. While the structures of RecA* and LexA have been solved separately, these efforts have led to vastly different models of complex formation, as evidenced by the suggestion that anywhere from three to seven RecA monomers might be involved in the protein interface^41–44^. Prior low-resolution negative stain electron microscopy studies helped to identify that LexA binds in the helical groove of the RecA* filament^43^. A more recent cryo-electron microscopy (cryo-EM) structure has shed some light on the interactions responsible for the SOS signal complex formation^45^; however, this structure was determined using only the isolated LexA-CTD rather than full-length LexA, and binding in a manner incompatible with native full-length LexA. Thus, fundamental questions remain regarding the orientation of complex components, stoichiometry of interfacial interactions, and the obligate molecular contacts controlling LexA cleavage.

Here, we offer the first structural model of the complete SOS signal complex with full-length LexA and present biochemical evidence that supports key mechanistic interpretations of our model. We describe a surprising interface of the NTD with RecA* that rationalizes exclusion of DNA-bound LexA and the resulting orchestration of the SOS response chronology. We further demonstrate that the LexA binding interface, comprised of three RecA monomers, a single activated RecA monomer containing the appropriate ‘key’ residue is sufficient to allosterically-induce LexA autoproteolysis. Taken together, our structure offers a refined mechanistic understanding of the DNA damage response and will inform efforts to target the adaptive evolution of antibiotic resistance.

## RESULTS

### Complete Structure of the SOS Complex

To assemble the ternary SOS complex, we prepared RecA* filaments with ATPγS and a 90-mer ssDNA template, followed by incubation with full-length catalytically-inactive LexA (K156A) to decorate the filaments under equilibrium binding conditions. After vitrification and imaging with cryo-EM, we employed helical refinement to reconstruct RecA* filaments. Unlike prior structural modeling that employed CTD-only LexA, here we found that full-length LexA is not uniformly bound throughout the RecA* filament groove. To overcome particle stack heterogeneity, we utilized 3D classification to parse LexA-bound and LexA-unbound sections of the filament, using a mask to address out-of-register binding (**Fig. S1**). From 1.77 million total initial particles, we ultimately obtained 233,920 sorted particles with good orientation distribution for further refinement, and subsequently produced a 2.93 Å resolution map focusing on LexA bound to the RecA* filament (**Fig. 1B, S1, S2A, S2B Table 1**).

**Table 1:** Cryo-EM data collection, refinement, and validation statistics.

|  |  |
| --- | --- |
| **EM Data Collection** |  |
| Facility | University of Pennsylvania |
| Microscope | Titan Krios |
| Camera | Summit K3 |
| Acquisition | EPU |
| Magnification | 64,000 |
| Voltage (keV) | 300 |
| Defocus range ($\mu\text{m}$) | -0.5 to -2.0 |
| Pixel size ($\text{\AA}$) | 1.38 |
| Electron exposure ($\text{e}^-/\text{\AA}^2$) | 46.2 |
| Micrographs (#) | 2,713 |
| Initial particles (#) | 1,770,386 |
| **Map** |  |
| Symmetry | C1 |
| Final particles (#) | 233,920 |
| Map resolution ($\text{\AA}$) | 2.93 (3.35) |
| FSC threshold 0.143 (0.5) |  |
| Resolution range ($\text{\AA}$; min/max/median) | 2.83/10.35/4.02 |
| Helical rise ($\text{\AA}$) | 16.2 |
| Helical twist ($^\circ$) | 59.2 |
| **Refinement** |  |
| Initial models used | AlphaFold2-generated |
| Model resolution ($\text{\AA}$) | 1.67 (3.03) |
| FSC threshold 0.143 (0.5) |  |
| Sharpening $B$ factor ($\text{\AA}^2$) | -70.5 |
| **Model composition** |  |
| Non-hydrogen atoms (#) | 22,876 |
| Residues (#) | Protein (2914) Nucleotide (27) |
| Ligands (#) | ATPgS (8) Mg (8) |
| **Correlation model vs. data** |  |
| CC (mask, peaks, volume) | 0.79, 0.58, 0.80 |
| CC (ligands) | 0.82 |
| **R.m.s. deviations** |  |
| Bond lengths $\text{\AA}$ (# > $4\sigma$) | 0.006 (0) |
| Bond angles $^\circ$ (# > $4\sigma$) | 0.752 (0) |
| **Validation** |  |
| MolProbity score | 1.62 |
| Clash score | 6.59 |
| EMRinger score | 3.17 |
| Rotamer outliers (%) | 0.78 |
| **Ramachandran plot** |  |
| Favored (%) | 96.20 |
| Allowed (%) | 3.80 |
| Disallowed (%) | 0.00 |
| **ADP (B-factors)** |  |
| Protein ($\text{\AA}^2$; min/max/mean) | 17.25/166.53/62.71 |
| Ligands ($\text{\AA}^2$; min/max/mean) | 10.29/63.37/39.23 |
| Nucleotides ($\text{\AA}^2$; min/max/mean) | 16.73/58.77/32.91 |
| **Accession codes** |  |
| EMDB | EMD-41579 |
| PDB | 8TRG |

In our refined structural model, the RecA* filament is organized in a right-handed, head-to-tail filament with a helical rise of 16.2 Å and twist of 59.2° (98.5 Å pitch with 6.08 RecA units per turn), in agreement with prior solved structures of RecA*^43,46–48^. At the interface of each RecA protomer, our high-resolution model shows clear density for both the bound ATPγS and the coordinated Mg^2+^ (**Fig. S2C**). At the helical axis, our model reveals the bound ssDNA stretched with nucleotide triplets maintaining B-form characteristics (**Fig. S2D**). Within the helical groove of RecA*, our model readily visualizes a complete ‘bound’ monomer of LexA, including the NTD and CTD, along with the CTD of the ‘unbound’ LexA monomer (**Fig. 1B**). Notably, the CTDs from each monomer of the LexA dimer are in different conformations. The RecA* ‘bound’ CTD occupies a cleavable conformation where active site Ser and the cleavage loop of LexA are closely approximated; the ‘unbound’ CTD, conversely, shows lower local structural resolution suggesting a higher degree of local flexibility, with the loop containing the scissile bond displaced from its associated protease active site (**Fig. 1C, S2A**). The NTD of the ‘unbound’ LexA monomer does not exhibit detectible density in our model, consistent with its flexibility and the dominant role of the CTD in enforcing dimeric LexA structure^26,49^.

The engagement of one complete monomer of the LexA dimer stands in stark contrast to prior proposed biochemical models where both CTDs were posited to bind across as many seven RecA monomers and to the recent structure with LexA-CTD only, where the role of the NTD remained unknown. In our complex, RecA* and the bound LexA monomer have extensive van der Waals contacts along the entirety of the monomer and within the nucleoprotein filament groove, along with several potential charge-charge interaction sites (**Fig. 1B, S5**). The LexA monomer binding spans a total of three RecA units in the RecA* complex (denoted as RecA1 to RecA3), with residues from each of the RecA L2 loop regions (res. 193-212) providing many of the contacts (**Fig. 2**). In our model, the L2 loops of the flanking RecA units (RecA1 and RecA3) appear to ‘brace’ the CTD and NTD of LexA, respectively, and the L2 loop of the middle RecA2 unit is engaged extensively with LexA (**Fig. 2**). The F203 residue in this “on-register” L2 loop is buried within a deep hydrophobic allosteric binding pocket that is formed in the closed conformation of LexA (**Fig. 2**). Comparisons indicate that the L2 loop from the on-register monomer (RecA2), and F203 specifically, is well defined, while the loop is much more flexible in the “off-register,” bracing RecA monomers (RecA1 and RecA3), with a loss of defined density for M202 and F203 in RecA1 (**Fig. 2**).

**Figure 2.**
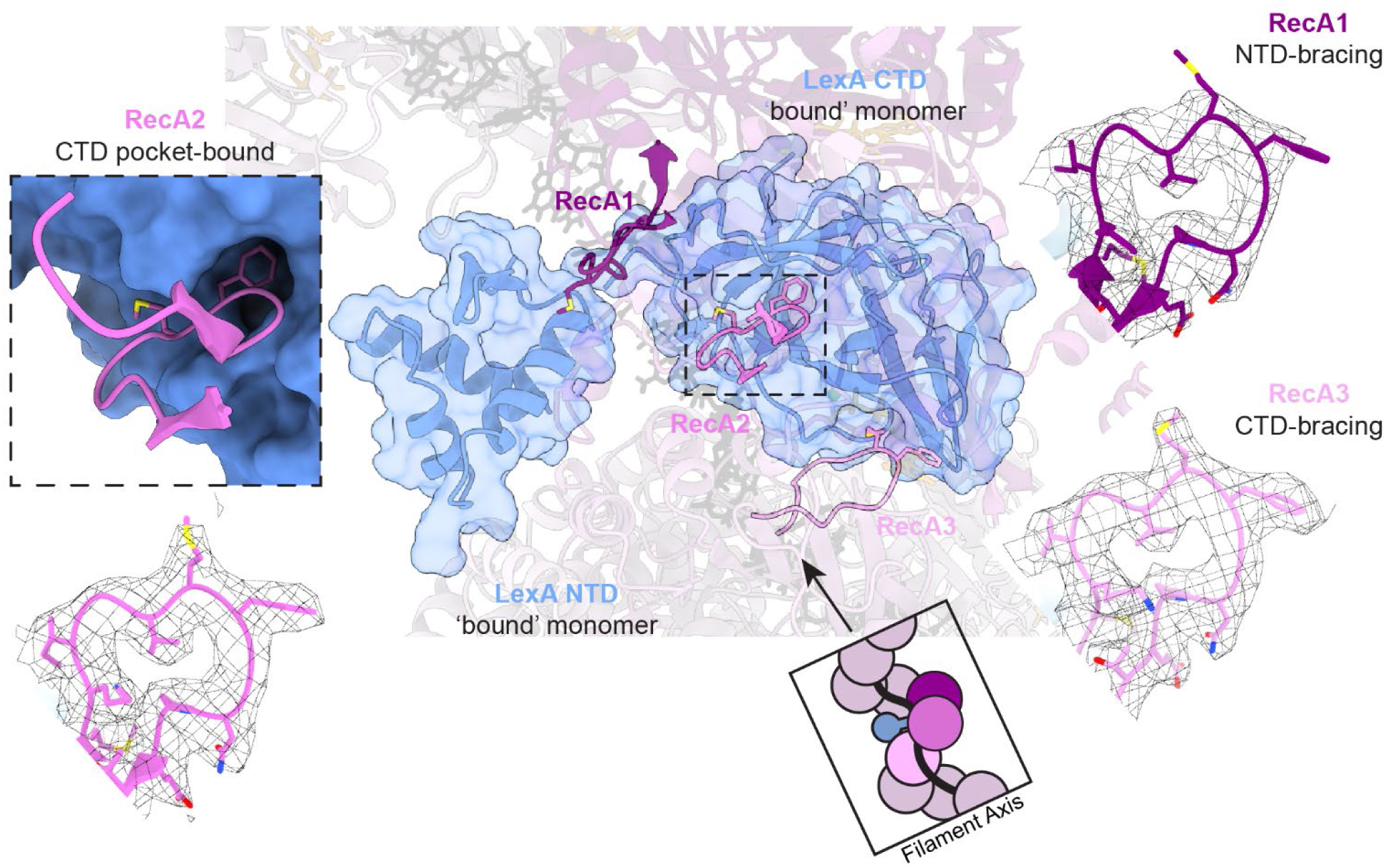
The role of the RecA L2 loops in complex formation. **A)** L2 loop regions (residues 197-212) of three successive RecA units are shown. At top left, the cartoon inset highlights the filament axis. At bottom left is a zoom in of the L2 loop from RecA2, with F203 binding in a deep allosteric pocket within LexA, defined by the residues shown at the bottom right. **B)** The electron densities for the L2 loops of the bracing RecA1 (top; purple), CTD-bound RecA2 (middle; magenta), and RecA3 (bottom; pink) are shown.

Given the structural insights suggested by our model of the complete SOS signal complex, we next aimed to refine our mechanistic understanding of RecA*-dependent LexA autoproteolysis, focusing attention on the role of the LexA NTD, the sufficiency of specific contacts for complex formation, and the importance of F203 engagement with LexA in SOS activation.

### The LexA NTD has a role in substrate discrimination

There are two different LexA species that could be viable substrates for RecA*- dependent cleavage in cells: DNA operator-bound LexA performing its repressor role or free cytosolic LexA. Prior work has suggested that early and late SOS pathway genes can be differentiated by the relative thermodynamic affinities of their operator sequences for LexA^34^. More detailed kinetic characterization has suggested that the specific timing for gene de-repression depends most on dissociation rates from DNA operator sequences, with slow off-rates observed for LexA from pro-mutagenic, late SOS pathway genes^34^. This latter study suggests a kinetic model in which the specific depletion of cytosolic LexA, but not DNA-bound LexA, leads to the sequential de-repression of genes. In this model, preferential interaction of RecA* with free LexA is assumed, otherwise RecA*-mediated cleavage of operator-bound LexA would disrupt the sequential de-repression of SOS pathway genes, which may prove transcriptionally costly and potentially lethal. However, the mechanism that could allow discrimination between these two LexA species remains unclear, and previous *in vitro* studies suggest that much of the LexA NTD is dispensable for both RecA* binding as well as cleavage stimulated by RecA* ^17,49,50^.

A striking attribute of our structural model of the SOS signal complex was the manner in which the LexA NTD engaged with RecA*. We considered the possibility that the ability of the LexA NTD to come into close proximity with the RecA* helical groove could serve a discriminatory function between DNA-bound and free LexA, since we would not expect any bulkier macromolecules tethered to the NTD to be able to occupy the same space. To explore this hypothesis, we examined the impact of steric effects in two different manners.

We first tested the ability of chimeric LexA constructs to be cleaved by RecA where the NTD was replaced with a more sterically prohibitive Maltose-Binding Protein (MBP) domain. In order to test proximity effects, we designed a series of chimeric catalytically active LexA constructs, where the CTD of LexA was tethered to either the native NTD (12 kDa) of LexA or with the globular MBP (42 kDa) via either the native short *E. coli* LexA linker (five amino acids) or the longer one modeled from *M. tuberculosis* (*Mtb*; 24 amino acids) (**Fig. 3A**). We additionally included a LexA-CTD only construct (FlAsH-LexA) where amino acids 1-74 have been replaced with a short peptide tag for fluorescent labeling (**Fig. 3A**)^50–52^. Notably, LexA auto-proteolysis can be induced by either incubation with RecA* or in a RecA* independent fashion by using alkaline conditions. To first assess any alterations to intrinsic cleavage activity, we compared rates of self-cleavage under alkaline conditions and found that all constructs cleaved at rates similar to wild-type LexA, indicating minimal impact of the NTD or linker on CTD conformational dynamics or cleavage (**Fig. S3**). By contrast, while the truncated FlAsH-LexA construct cleaves similarly to WT, the introduction of MBP in lieu of the native NTD markedly prevented RecA*-dependent cleavage (**Fig. 3A**). The introduction of the longer linker from *Mtb* leads to a partial rescue in cleavage, suggesting that spatial separation can reduce the steric blockage observed with MBP (**Fig. 3A**).

**Figure 3.**
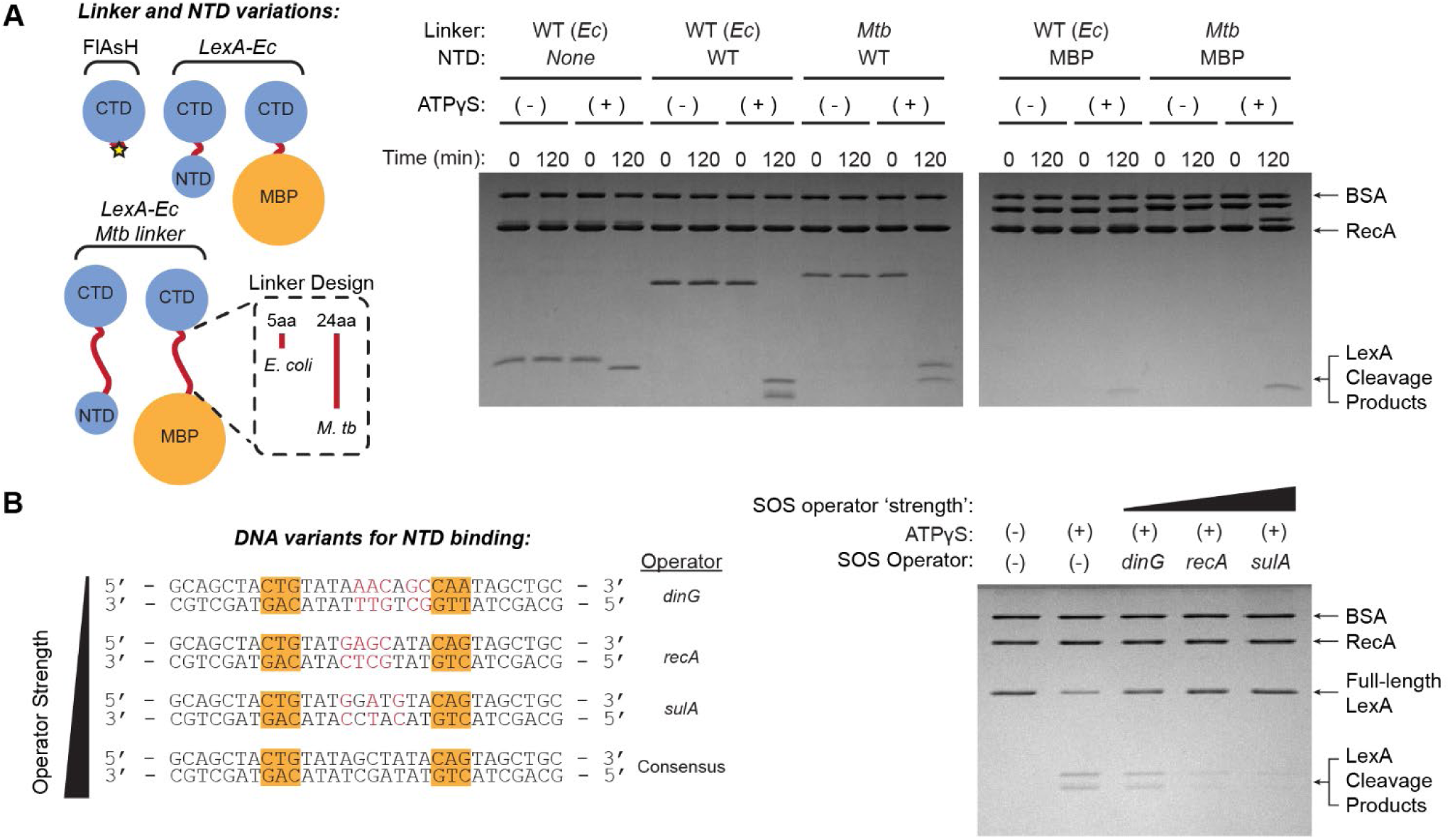
The role of the LexA NTD in RecA*-dependent proteolysis. **A)** At left, design of chimeric LexA constructs that include variations in the NTD (native, removed or replaced by MBP) and the linker (native or elongated *M.tb* linker). At right, SDS-PAGE analysis of RecA*-dependent cleavage of LexA variants. **B)** SOS operator design. At left are shown a consensus DNA operator sequence (at bottom), with the invariable region that defines the SOS box highlighted in orange. The *dinG*, *recA*, and *sulA* sequences are ordered in increasing strength and derived from their native promoters, with deviations from the consensus sequence highlighted in red. At right, SDS-PAGE analysis of RecA*-dependent cleavage of LexA with or without pre-binding of the SOS operator DNA.

Second, we tested an alternative and more physiologically relevant way to introduce steric bulk associated with the NTD by exploring cleavage when WT LexA was pre-incubated with excess operator DNA. We selected 40 base pair (bp) sequences embedding three different 20 bp SOS operators with distinct LexA binding kinetics – *dinG* (early induction; fast off-rate), *recA* (intermediate induction, intermediate off-rate), and *sulA* (late induction; slow off-rate) (**Fig. 3B**)^34^. We found that pre-incubation with these operators showed a distinct difference in qualitative RecA*-dependent cleavage, with the intermediate and late induction operators (*recA* and *sulA*) more potent at protecting LexA from cleavage *in vitro* (**Fig. 3B**). To examine the mechanisms at play, we generated a 20 bp tight-binding ‘consensus’ operator or an analogous 40 bp operator. Using these operators, we observed that DNA binding does not impact the intrinsic CTD dynamics or activity, with similar RecA*-independent cleavage rates at pH 7.5 in the presence or absence of operator DNA (**Fig. S4A**). To specifically probe binding of LexA to RecA*, we employed a variant of LexA containing the fluorescent unnatural amino acid, acridonylalanine (δ), which allows for tracking by fluorescence anisotropy ^49,53^. Using this assay approach, DNA-binding and RecA* binding both result in expected changes in anisotropy with LexA-δ. However, the presence of the 40 bp consensus operator leads to an intermediate anisotropy value, likely due to a mixed population of RecA*-bound LexA, and operator-bound LexA that is prevented from binding RecA*, resulting in an apparent decrease in overall anisotropy (**Fig. S4B**). Interestingly, although the LexA variant with the longer *Mtb* linker binds the consensus operator with similar affinity to native LexA, the longer linker results in some detectable rescue of RecA*-dependent LexA cleavage (**Fig. S4C, S4D**).

Taken together, our data indicate that although the NTD is entirely dispensable for RecA*-dependent LexA cleavage, the NTD must be sterically accommodated to allow for LexA positioning within the RecA* groove, with either unnatural steric bulk or the more physiological binding of DNA impeding complex formation. Thus, for the *E. coli* system, the NTD plays a critical role in the selective depletion of DNA-unbound LexA, offering a mechanistic explanation for control over the timing of SOS induced genes.

### RecA F203 engages an allosteric pocket to control LexA cleavage

We next explored molecular contacts spanning the LexA/RecA* interface to understand the determinants of LexA binding and induced cleavage. In addition to numerous van der Waals contacts, the interface exhibits several distinct patches of contacts with either long-range electrostatic interactions or potential hydrogen bonds. From the LexA perspective, these patches can be separated into a CTD patch, L2 loop stabilizing patch, and NTD patch (**Fig. S5A-C**). Given our prior assessment of the role of the NTD, we focused analysis on the L2 stabilizing patch and the CTD patch and systematically substituted candidate residues with alanine. In this screen, most charged contacts showed no significant effect on RecA*-independent cleavage and were individually dispensable for RecA-dependent cleavage (**Fig. S3, S5D, S5E**). Although impacts were modest, the largest effects were seen with LexA residues E95 and D151 from the CTD patch and residues K135 and E189, which hydrogen bond with the RecA L2 loop backbone (**Fig. S5**). Notably, while in prior work with the CTD-only complex, groups of charge reversal mutations were suggested to greatly impact LexA proteolysis, our alanine scan suggests that these contacts to be individually of lesser importance to the overall interaction^45^.

As noted earlier, the L2 loops form a substantial portion of the interface, with bracing interactions in off-register RecA1 and RecA3 monomers and more extensive engagement with on-register RecA2. Within the RecA L2 loop, residue F203 is deeply buried in a hydrophobic pocket on LexA (formed by residues V79, V82, F111, L113, V146, and V153) that exists exclusively when LexA is in the ‘cleavable’ conformation (**Fig. S6A**). This allosteric pocket is on the opposite face of the LexA active site, with the L2 loop positioning F203 for insertion into the pocket. This insertion appears to stabilize ordered beta-strand formation from the otherwise disordered loop in LexA containing the scissile bond.

In prior work on LexA, the ‘cleavable’ state was originally solved via X-ray crystallography using a LexA quad-mutant (QM) consisting of L89P, Q92W, E152A, and K156A (PDB: 1JHE)^29^. The K156A mutation renders LexA incapable of autoproteolysis, while the L89P, Q92W, and E152A mutations were shown to stabilize the cleavable conformation. Comparing the QM crystal structure with our cryo-EM model showed an unexpected and striking correlation. In the QM structure, the Q92W mutation results in a significant reorientation of residue 92, which normally sits at the brim of the allosteric pocket, such that the six-membered ring of the indole group is now situated nearly identically to the RecA F203 phenyl group and positioned into a newly formed allosteric pocket (**Fig. 4A**).

**Figure 4.**
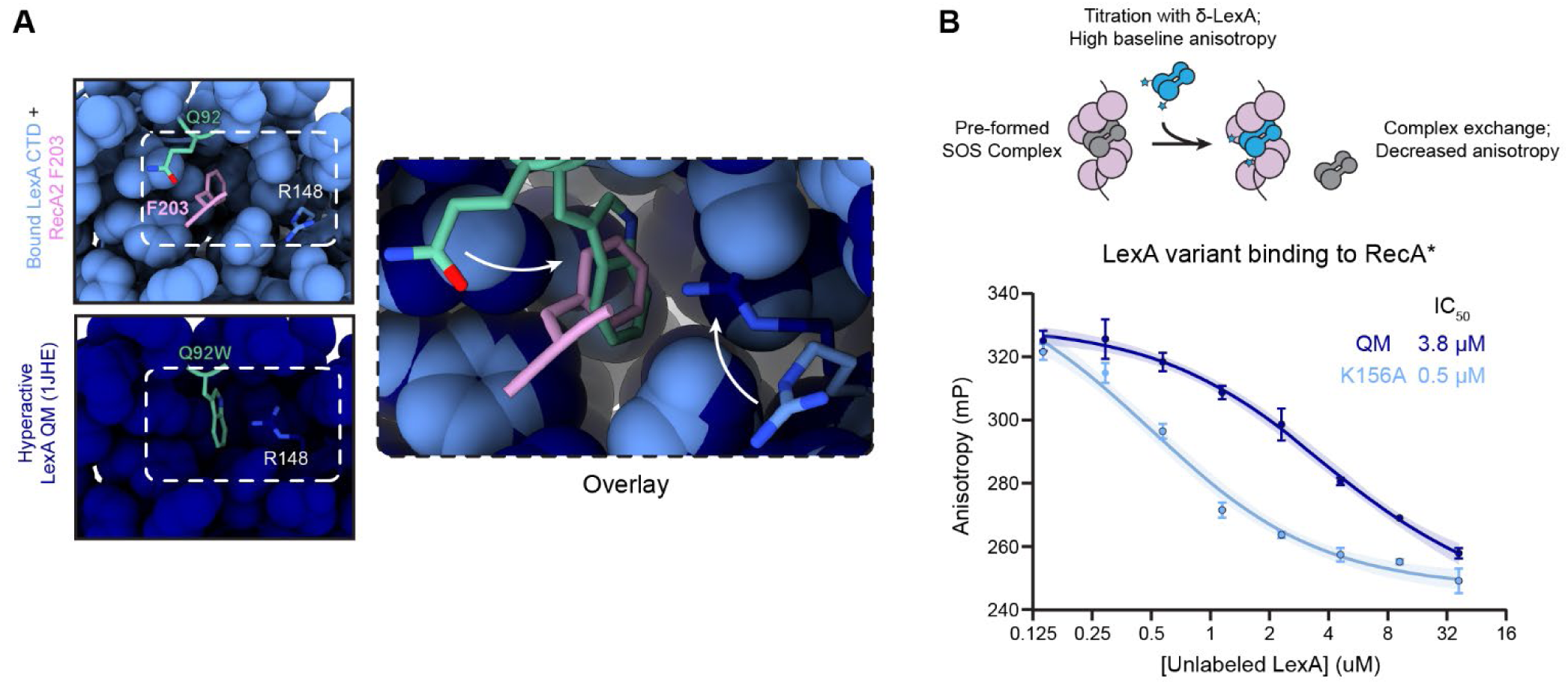
The hyperactive LexA variant auto-populates the allosteric pocket in a similar manner to RecA*-bound LexA. **A)** Comparison between the RecA*-bound LexA model in our presented structure (light blue) and the crystal structure of the hyperactive LexA QM (1JHE, dark blue). Residue 92, which is a Gln in WT LexA and Trp in the QM, is highlighted in light green. The overlay demonstrates the reorientation to residue 92 to permit auto-population of the allosteric pocket. **B)** Binding of LexA QM or LexA K156A to RecA* as measured by competition against LexA-δ. Cartoon schematic for the assay design is shown above graph. Data shows the mean from three replicates and error bars indicate standard deviation. Data are fit against a dose response curve (three parameters) with shaded regions showing 95% confidence intervals and best fits for IC_50_ given.

We reasoned that if F203 binding within the allosteric pocket is a critical determinant of LexA engagement, then RecA* filaments should be less capable of binding the LexA QM which auto-populates this pocket. To test this hypothesis, we employed our highly sensitive LexA-δ binding assay^49^. Here, the SOS signal complex is first pre-formed with RecA* filaments and a LexA variant of choice. LexA-δ is then titrated into the complex with competitive binding tracked by fluorescence anisotropy. Using this approach, we observed that the LexA QM has nearly 8-fold weaker binding relative to LexA K156A (**Fig. 4B**) supporting the hypothesis that RecA F203 insertion is a key determinant of LexA binding.

The RecA L2 loop has been previously characterized for its role in filamentation and homologous recombination. For example, a past study by Hörtnagel et al. performed saturation mutagenesis of the L2 loop, including F203, to explore essential residues for recombinational repair in *E. coli*^54^. Given the multiple roles of RecA, we recognized that the *in vivo* requirements might be more stringent than the requirements for SOS activation. We therefore expressed and purified a subset of the F203 mutants, including a mutant where propargyl-tryosine (ppY) was incorporated via amber suppression, adding steric bulk at the *para* position. Each mutant was assayed for its ability to 1) form RecA* filaments via fluorescent anisotropy changes upon binding to a fluorophore-labelled ssDNA, 2) bind to LexA using the LexA-δ binding assay, and 3) stimulate LexA cleavage *in vitro*. In our filamentation assay, all tested substitutions had either no statistically significant effect (Y, S, A, M, I, Q, E) or a small negative effect (ppY, W) relative to RecA WT (**Fig. 5A**). Despite proficiency in filamenting on ssDNA, however, only a subset of the mutants was capable of binding to LexA (ppY, Y, W, M, I) to varying degrees (**Fig. 5A**). Given that the polar and charged variants were not tolerated, the LexA binding activity tracked with our hypothesis about the critical role for van der Waals interactions in the key allosteric pocket. Strikingly, in our cleavage assays, the degree of binding impairment correlates with the cleavage activity impairment, suggesting both that F203 is a major determinant of SOS complex formation and that LexA binding is, in turn, the determinant of the proficiency of cleavage (**Fig. 5A, B**).

**Figure 5.**
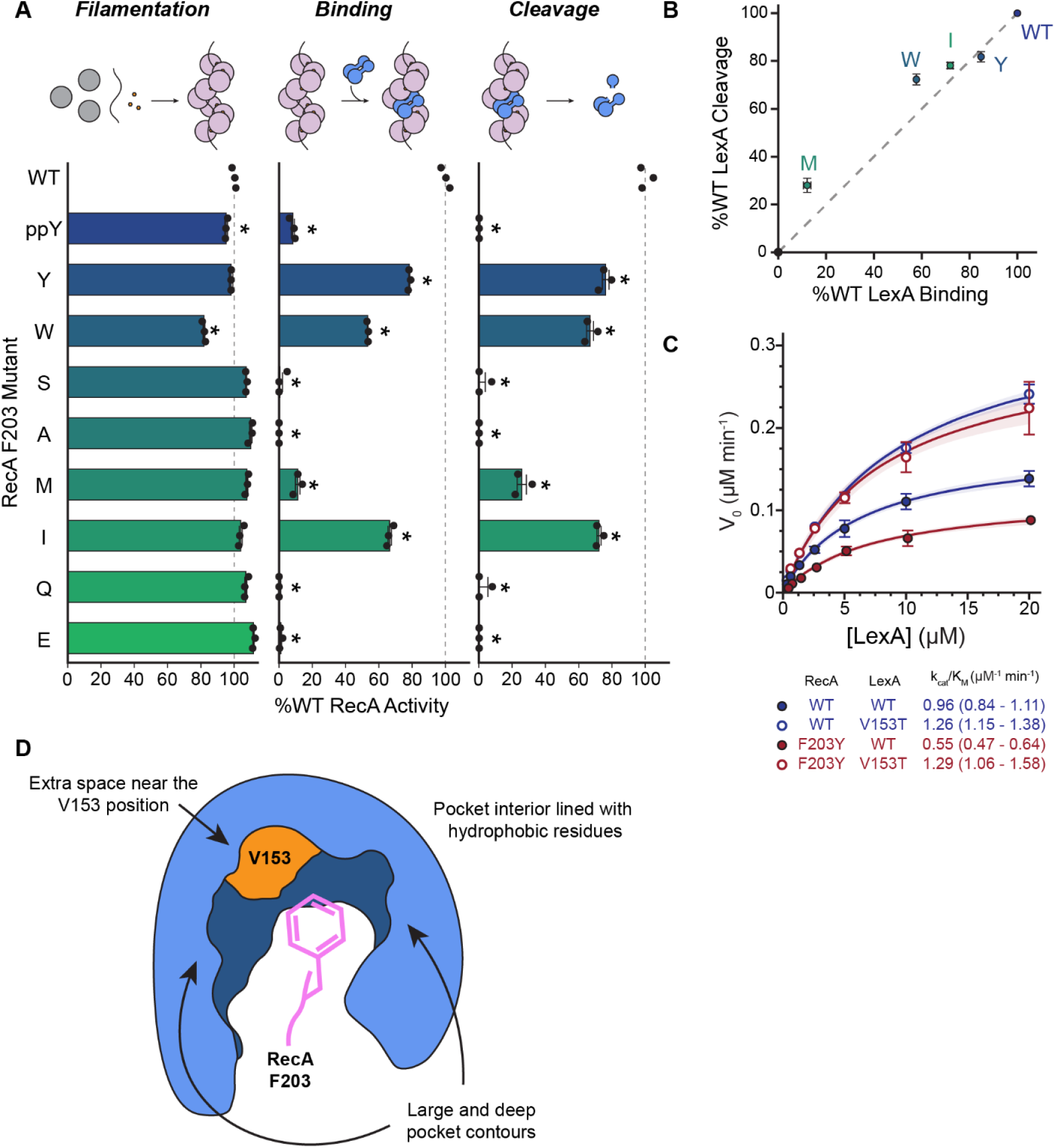
RecA F203 variants modulate binding and dictate LexA cleavage efficiency. A) RecA F203 mutant filamentation, binding, and cleavage expressed as a percentage of RecA WT anisotropy (normalized to 100% shown by the dotted line, defined from the average of the three WT points) in the FAM-ssDNA binding assay, LexA-δ binding assay, and LexA cleavage assay, respectively. Data show the means of three replicates, with error bars denoting standard deviation. * denotes significance, with a greater than 95% posterior likelihood of being less than RecA WT. **B)** Plot showing the %WT LexA Binding versus %WT LexA cleavage for the active F203 mutants. Data show the means of three replicates, with error bars denoting standard deviation. **C)** Steady-state kinetics of orthogonal RecA/LexA pairs are shown. Plots of initial velocities of LexA cleavage as a function of [LexA], measured by the quantification of LexA cleavage product in the sensitive fluorescent SDS-PAGE LexA cleavage assay from three replicates. Error bars denote the standard deviation. Solid lines show the best fit to a modified Michaelis-Menten equation, with 95% confidence intervals shown (shaded regions). The best fit values of the specificity constants are noted, with 95% confidence intervals shown in parentheses. **D)** Cartoon representation of the F203-binding pocket, highlighting different pocket features.

In line with our model, phylogenetic analysis reveals that this F203 position is highly conserved, along with many of the hydrophobic pocket residues on LexA, with the exception of V153 which is more variable (**Fig. S6B**). We noted the occasional presence of Y203, M203, and W203 variants across species, including *T. thermophilus* where Y203 is paired with T153 in its cognate LexA, with the potential for a hydrogen bond within the pocket. To explore this evolutionary divergence, we engineered RecA F203Y and LexA V153T mutations and analyzed cleavage of these variants under steady-state conditions. Exploring the permutations of LexA and RecA, we observe that although V153T alone is tolerated, RecA F203Y results in a 2-fold reduced specificity constant for LexA WT co-protease activity (**Fig. 5C**). In line with our prediction, however, when the F203Y mutation is paired with V153T it demonstrates a clear rescue phenotype relative to F203Y alone (**Fig. 5C**), highlighting a degree of plasticity across species in this critical LexA-RecA* interface.

### A single RecA F203 is sufficient for SOS activation

One major reason that the SOS complex has been resistant to biochemical characterization is that mutations in one RecA monomer propagate through every monomer of the filament, leaving ambiguity as to the specific contacts responsible for SOS activation. In our model, this issue surfaces with the critical F203 interaction, as impacts on LexA binding/cleavage impairment could be attributed either exclusively to effects on the CTD-pocket bound L2 loop (RecA2), or to compounded effects from losing F203 in the NTD-bracing and CTD-bracing loops as well (RecA1 and RecA3).

To resolve this ambiguity and determine the critical interface for SOS activation, we explored an engineered oligomeric RecA construct, RecA3x, containing three covalently tethered RecA units mutated on the end monomers to prevent filament extension (**Fig. 6A**). We recently demonstrated RecA3x as the ‘minimal’ RecA* species competent for LexA cleavage, as an analogous RecA2x construct was unable to make stable filaments and to support LexA cleavage^44^. Using the RecA3x construct, individual RecA monomers can be mutated either in isolation or in all combinations (**Fig. 6A**). As anticipated, based on the monomeric RecA F203A mutant, we found that all permutations of one, two or three mutant monomers were equivalently capable of stable filament formation relative to unmutated RecA3x (**Fig. S7A**). Moving to LexA cleavage, when F203A is mutated in all three RecA3x protomers, LexA cleavage is abolished as anticipated (**Fig. 6B**), a result aligning with the inability for this variant to bind to LexA (**Fig. S7B**). By contrast, when F203A is only present in any one or two of the monomers, we observed that binding to LexA and co-protease activity are observable indicating that a single native F203 is sufficient to support SOS activation (**Fig. 6B, S7B**). A closer look at the pattern of cleavage helps to provide a more comprehensive view. Although competent for LexA cleavage, the variant with an F203A mutation in the latter two RecA protomers was most severely impacted, and, correspondingly, mutation to F203A in the first protomer alone was the least effected construct for both LexA cleavage and binding (**Fig. 6B, S7B**). This corresponds to the number of maintained contacts, as expressed by the calculated surface area between possible RecA/LexA contact regions based on the position of the “docking” monomer containing the unmutated F203 (**Fig. 6B**). These results support a model where, although three RecA monomers are needed to make a stable RecA* filament, a single RecA monomer in the active ATP-and DNA-bound conformation is sufficient for LexA binding and co-protease activity. While sufficient, the added bracing contacts with an upstream RecA monomer can further stabilize the complex to support more efficient LexA cleavage. Thus, conceptually, the ‘minimal’ complete SOS complex involves LexA bound to activated RecA3x containing a single F203 in the central monomer.

**Figure 6.**
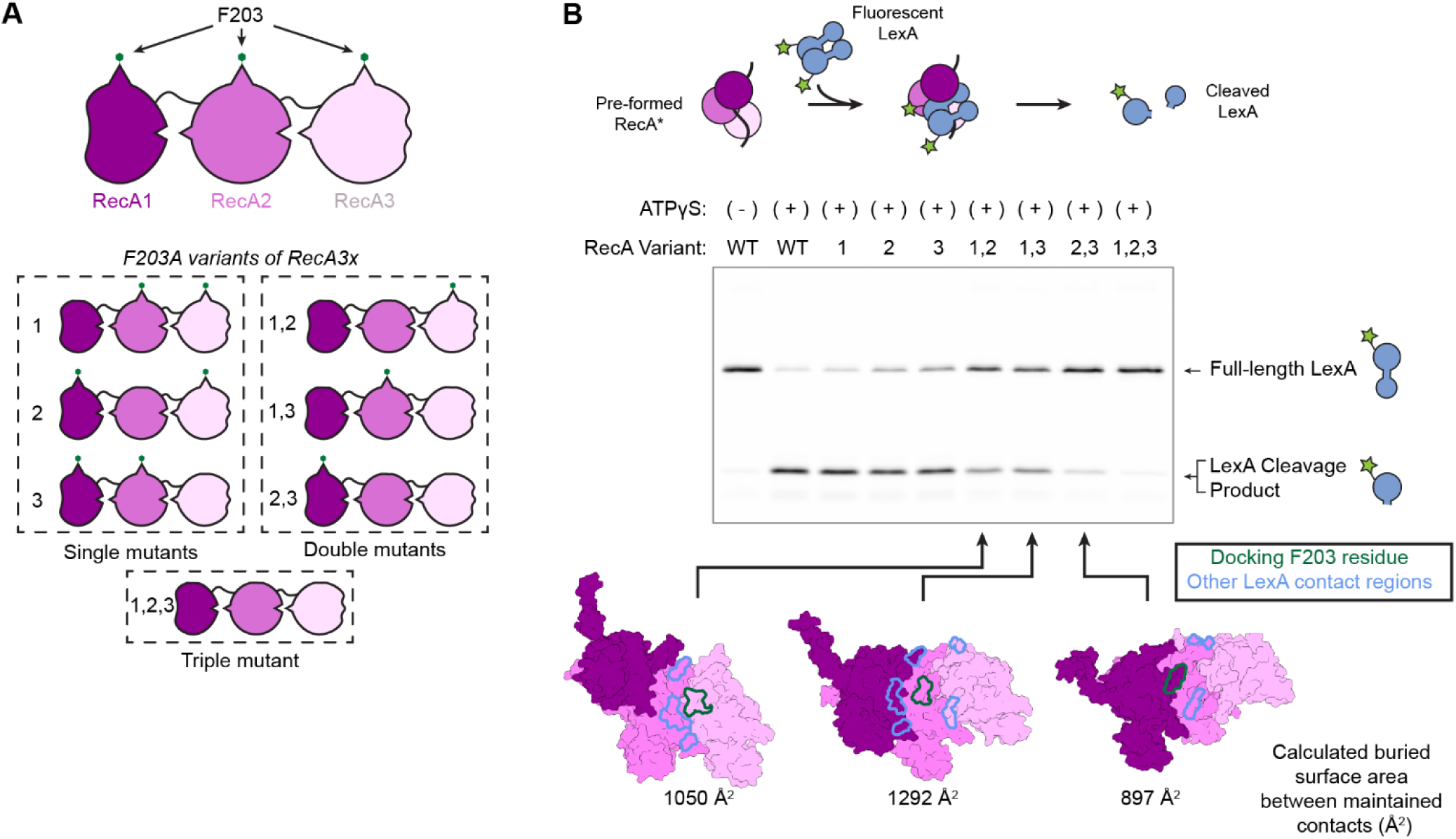
A single active monomer with F203 in RecA* is sufficient for LexA cleavage. **A)** The RecA3x construct involves covalent linkages of three RecA monomers, allowing for mutations to be made in individual monomers. Below are shown the permutations of single, double and triple mutants evaluated in the RecA3x scaffold. **B)** RecA3x cleavage assay. The schematic describes that RecA3x variants were used to performed an activated RecA* and then incubated with fluorescent LexA-CF, whose cleavage can be analyzed by SDS-PAGE as shown below. Focusing on the three double mutants, at the bottom are shown surface models of RecA trimer in the active conformation, with the putative “docking” L2 loop (green) and the remaining available contact regions with LexA (blue). The numbers below correspond to the calculated buried surface area available between RecA* and LexA for each register of LexA binding.

## DISCUSSION

Although the SOS complex has been a focus of study for nearly half a century, a high-quality atomic-resolution model of the RecA*-LexA complex has remained elusive. Extensive mutagenesis studies have failed to elicit LexA mutants that are specifically deficient in RecA*-induced cleavage, but not autoproteolysis. Similarly, mutations that specifically prevent LexA cleavage, without impacting other RecA functions have been difficult to assimilate into a functional model, as mutations in one monomer propagate to every monomer in RecA*.

Against this backdrop, resolution of the complete SOS complex structure offers the possibility of insights into both the underlying biological mechanisms regulating SOS activation and opportunities for the design of antagonists. A recently published model helped to reveal some of the structural features of the SOS complex^45^. However, the generation of this model required that the macromolecular complex be constrained to a non-physiologic form: a RecA* filament fully decorated with a truncated form of LexA repressor lacking the DNA binding NTD. To overcome these limitations, we instead purified a macromolecular complex more representative of the biologically relevant species and exploited 3D classification, masking, and local refinement to resolve heterogeneity in full-length LexA occupancy of the otherwise helically symmetric RecA* filament.

By capturing the LexA NTD in our model of the RecA*-LexA complex, we provide structural evidence that the NTD positions itself within the helical groove of RecA* in a ‘side-on’ configuration involving both domains of LexA. On one hand, the finding that the LexA NTD is closely associated with the RecA* helical groove was surprising in that the NTD is entirely non-essential for RecA*-dependent cleavage^45,49,50,52^. On the other hand, the observation provides a rationale for the ordered timing of SOS response genes, a key attribute of the DNA damage response. The close association of the NTD suggested a role in steric discrimination, as we find that either replacement of the NTD with a bulky MBP or binding to operator DNA impedes both RecA* binding and LexA cleavage. Steric discrimination provides a mechanism for specific depletion of LexA from cytosolic pools in *E. coli*. This binding discrimination explains how differences in dissociation rates of LexA from its various operators can control the differential timing of early DNA repair versus late genes, including genes that could have adverse consequences if induced early, such as the gene products halting bacterial replication and septation^34,55^. Our observations may also hold implications for the differential regulation of the SOS response in other bacterial species. Although the inter-domain linkers in most bacterial pathogens are short, as in *E. coli*, the linker can be > 60 amino acids in some species. As cleavage inhibition was partially alleviated by substituting the longer *Mtb* linker into *E. coli* LexA, it is feasible that cleavage could occur on both cytosolic and DNA-bound LexA in some species, with resultant changes in the chronology or stochasticity of SOS genes’ induction.

In prior work, we identified an activated RecA trimer as being the minimum filament size required for LexA cleavage, a finding now corroborated by our structural model showing LexA chiefly engaged with RecA* along three successive protomers^44^. This model notably differs from that suggested by the recent structure with isolated LexA-CTD, where only two RecA monomers were modelled as part of the complex. While the central RecA monomer provides dominating contacts, we identified a ‘bracing’ role for the L2 loops on the off-register RecA monomers, with our RecA3x mutant series offering evidence for a greater importance to NTD over CTD bracing contacts (**Fig. S5**).

In the ‘on register’ RecA monomer, F203 from the L2 loop inserts into the critical allosteric pocket found at a distance from the active site and scissile bond. Much like a key inserting into a lock, the F203 residue inserts into this pocket found only in the cleavable variant of LexA, thereby stabilizing the cleavable conformation. While all F203 variants were able to make RecA* filaments in our mutational studies, substitutions in this essential residue resulted in altered LexA binding, which then directly correlates with altered LexA cleavage kinetics. Interestingly, this observation could help explain previously reported patterns for *in vivo* tolerance to substitutions at position F203^54^, as F203 was relatively tolerant to mutation in a recombination plaque-based assay, but more restrictive under UV challenge, where both recombination and SOS activation are thought to be of importance.^54^ The dual roles of RecA highlight competing evolutionary pressures, where inducible SOS activation could be a determining factor for the general conservation observed at F203 in RecA.

The allosteric pocket in LexA also offers a compelling target for inhibition. Small molecule antagonists could either prevent the formation of this allosteric pocket or occupy it in a manner that excludes F203 and RecA* while preventing LexA cleavage. This deep and well-defined pocket provides a uniquely accessible binding interface for rational inhibitor design in comparison with typical protein-protein interaction interfaces which tend to be shallower and cover a broader surface. It is notable that while many of the pocket residues have small, aliphatic sidechains, there is significant variation of the specific residues in the pocket. While offering a viable target, this evolutionary tolerance may also present some challenges in mutability whereby mutations within the pocket contour region could lead to exclusion of small molecules, or that existent species variation could limit the breadth of potential SOS antagonists. Beyond inhibition, one could also envision small molecules that could mimic allosteric activation and specifically activate the SOS response in a DNA damage-independent manner. Such agents would provide a convenient tool to decoupling the biological outcomes of SOS activation from recombinational repair in response to different cellular events.

Given the dire need for novel approaches to counteract bacterial pathogens, the complete absence of LexA homologues from mammalian biology makes targeting the SOS response an intriguing opportunity. We anticipate that the structural insights into the complete SOS complex provided here will significantly enhance efforts to develop SOS antagonists, an outcome that would benefit not only future studies into biologic stress response pathways but also therapeutic directions that target the evolution of antibiotic resistance.

## METHODS

### Recombinant Protein Cloning, Expression, and Purification

Constructs were cloned into pET41 expression vectors, harboring either an N-term poly-His tag, with the exception of unnatural amino acid containing constructs (LexA-δ, incorporated at residue 161, and RecA F203ppY) which contain a C-term poly-His tag. Engineered RecA3x constructs were designed as previously described^44^.

LexA constructs were expressed in BL21(DE3) cells whereas all RecA constructs were expressed in the *ΔrecA* BLR(DE3) cells (Thomas Scientific). The acridonylalanine (δ) labeled LexA construct, LexA Q161δ, was expressed in BL21(DE3) cells containing an auto-inducible plasmid-borne tRNA synthetase for δ in the presence of 1.0 mM solubilized δ in the media^56^. Overnight cultures were grown in MDAG non-inducing media and diluted 1:100 into fresh LB media. Cultures were grown to an OD600 of 0.6-0.8 (LexA) or 1.2-2.0 (RecA) before induction with 1 mM IPTG. After four hours at 37 °C, harvested cells were resuspended in either LexA lysis buffer (20 mM sodium phosphate, pH 6.9, 0.5 M NaCl, 25 mM imidazole) with cOmplete EDTA-free protease inhibitor cocktail tablet (Sigma) or RecA lysis buffer (50 mM sodium phosphate, pH 8.0, 500 mM KCl, 500 mM NaCl, 25 mM imidazole, 0.1% v/v Triton X-100, dissolved cOmplete EDTA-free protease inhibitor tablet) and stored at -80 °C.

Thawed cells were lysed by sonication with 250 µg/mL lysozyme and 25 units/mL benzonase. Lysates were clarified at 25,000 rcf for 30 min, and the supernatant was flowed over HisPur cobalt resin (Thermo) pre-equilibrated with the respective lysis buffer. The resin was washed with 20 column volumes (CV) of either LexA lysis buffer or RecA cobalt wash buffer (50 mM sodium phosphate, pH 8.0, 500 mM KCl, 500 mM NaCl, 100 mM imidazole), followed by elution in 3 CV of LexA cobalt elution buffer (20 mM sodium phosphate, pH 6.9, 500 mM NaCl, 400 mM imidazole) or RecA cobalt elution buffer (50 mM sodium phosphate, pH 8.0, 500 mM NaCl, 400 mM imidazole). For further purification, RecA samples were precipitated by incubation for 30 min at 4 °C using 48% (final) saturation ammonium sulfate, and the pellet was retained after spinning at 3,000 rcf for 30 minutes. Pellets were resuspended in RecA heparin load/wash buffer (20 mM Tris-HCl, pH 8.0, 20 mM NaCl, 2 mM TCEP) and dialyzed overnight at 4 °C against the same buffer to remove residual ammonium sulfate. LexA samples were diluted to <300 mM (final) NaCl using LexA heparin load/wash buffer (20 mM Tris-HCl, pH 7.0, 200 mM NaCl). All samples were filtered with a 0.2 µm GF+PES filter and loaded onto two in-line Heparin HiTrap HP 5 mL columns (Cytiva) using an ÄKTAPurifier system. The column was washed sequentially with 0%, 10%, 20%, and 30% mixes of wash buffer and the appropriate elution buffer (RecA: 20 mM Tris-HCl, pH 8.0, 1.5 M NaCl, 2 mM TCEP; LexA: 20 mM Tris-HCl, pH 7.0, 1.5 M NaCl) and all fractions were collected. Fractions were dialyzed overnight at 4 °C into RecA storage buffer (20 mM Tris-HCl, pH 8.0, 200 mM NaCl, 2 mM TCEP, 10% glycerol) or LexA storage buffer (50 mM Tris-HCl, pH 7.0, 200 mM NaCl, 0.5 mM EDTA, 10% glycerol). RecA fractions were tested for filamentation capacity using the filamentation assay (see below), and the most active fraction was retained. LexA fractions were checked for purity via SDS-PAGE, and pure fractions were pooled. All protein concentrations were determined using Qubit and aliquots were stored at -80 °C. A representative gel showcasing each major construct in this study is shown in **Fig. S8**.

The His-LexA K156A construct that was to be used for cryo-EM was further processed to remove the His-tag by incubation with thrombin protease overnight at 4 °C in thrombin cleavage buffer (50 mM Tris-HCl, pH 7.0, 200 mM NaCl, 10 mM CaCl2). The post-cleavage sample was then flowed over HisPur resin pre-equilibrated with LexA storage buffer, and flowthrough was collected and exchanged into fresh storage buffer to yield untagged LexA K156A.

### Cryo-EM grid preparation and data collection

RecA* was pre-formed by mixing 5 µM His-RecA WT with 1 µM [oligo] GGT 90mer ssDNA substrate (GGT90, GGT repeats of 90 nucleotides total length) in 1X cryo-EM activation buffer (70 mM Tris-HCl, pH 7.5, 150 mM NaCl, 1 mM MgCl2, 2 mM TCEP, 0.25 mM ATPγS) and incubated for 20 minutes at room temperature. LexA K156A was then spiked into the reaction to a final concentration of 25 µM (10-fold molar excess to RecA) and incubated for an additional 30 minutes at room temperature. 5 µL of reaction mixture was applied to glow-discharged Quantifoil R 1.2/1.3 copper 300-mesh holey carbon grids followed by blotting and plunge-freezing into liquid ethane on a Mark IV Vitrobot (7.0 s blot time under 100% humidity at 25 °C). Grids were stored in liquid nitrogen until loaded into the Titan Krios for data collection.

2,748 movies were collected on a Titan Krios microscope at 300 kV in nanoprobe mode, with a K3 Summit in counted super resolution mode. Magnification was set to 64,000 for a final pixel size of 0.69 Å and defocus range of -0.5 to -2.0 µm. Movies were collected with EPU software and fractionated over 40 frames with aberration-free image shift (one target per hole) for data acquisition. The exposure rate was 3.4-3.7 electrons per pixel per second for a total nominal exposure of 45.5 *e*/Å^2^.

### Cryo-EM Data Processing

Processing of micrographs was done using CryoSPARC v 4.0.3.^57^ All 2,748 movies were imported into CryoSPARC followed by Patch Motion Correction with a Fourier cropping factor of 0.5. Local CTF values were estimated with the Patch CTF Estimation. Maximum CTF fit resolution cutoffs of 2.77 Å and 20 Å were used to filter micrographs, with 2,713 micrographs selected for downstream processing. The automated filament tracer was used to select particles, using a filament diameter of 100 Å (90-150 Å) and a separation distance of 17 Å. This resulted in 3,423,408 total particles. These particles were then subjected to reference-free 2D classification to remove hole edges and aberrantly selected particles. Twelve 2D classes were chosen, with 1,770,386 total particles. The particle stack was asymmetrically refined using the Helical Refinement job in CryoSPARC with non-uniform refinement selected, optimizing for symmetry parameters. Using the optimized symmetry parameters, Helical Refinement was run again to obtain a resolution of 3.06 Å determined by half-map Fourier shell correlation (FSC) at 0.143.

To recover the LexA bound to RecA* in the groove, the 1,700,386 particles were subjected to 3D classification with a mask covering a space within the groove. A single class containing clear density of LexA in the groove, containing 360,867 particles, was then further classified with 3D classification into 3 classes using enforced hard classification. Two of the resulting 3D classes, corresponding to those with more defined CTD densities, were combined for a final particle stack of 233,920 particles.

This final particle stack was then taken through another round of Helical Refinement followed by Local Refinement for a final map resolution of 2.93 Å. This final map was sharpened using DeepEMHancer with a tightMask algorithm^58^. The final particle stack was analyzed for orientation distribution using cryoEF, with a calculated efficiency of 0.67 (**Fig. S2BC**)^59^. The final map was also assessed for local resolution using the local-resolution estimation in Relion version 3.1.3 (**Fig. S2A**)^60^.

### Model Building and Refinement

Initial models of each chain were generated using the AlphaFold2 plugin in Benchling and initially fit into the density using rigid body fitting in ChimeraX v 1.4^61,62^. Bound ssDNA was built manually in *Coot*, and ATPγS ligands were built and fit using Phenix eLBOW^63–65^. This initial model was then iteratively refined using Phenix’s real-space refinement, with manual adjustments in *Coot*^66^. Final refinements were done in ChimeraX using ISOLDE^67^. For model validation, we used Phenix’s comprehensive validation for cryo-EM models^68,69^. Model quality was assessed using a variety of parameters available in the comprehensive validation output, including MolProbity score, clashscore, Rama-Z score, bond length and angle outliers, and overall map-to-model agreement.

### LexA Alkaline Autoproteolysis Assays

LexA constructs were prepared by diluting 4X protein working stocks in nuclease-free water to a final [LexA] of 5.0 µM. These were then mixed 1:1 with alkaline LexA autoproteolysis buffer (100 mM Tris-Gly-CAPS, pH 10.6, 300 mM NaCl) and incubated in a 25 °C water bath. At each time point, 10 µL reaction mixture was added to 10 µL 2X Laemmli buffer and boiled at 95 °C for 5 minutes. 10 µL of each sample was then loaded onto a 15% SDS-PAGE gel for analysis.

### RecA-dependent LexA Cleavage Assays

Activated RecA* was prepared by mixing RecA and GGT 18mer ssDNA substrate together in 1X activation buffer (70 mM Tris-HCl, pH 7.5, 150 mM NaCl, 10 mM MgCl2, 2 mM TCEP, 0.25 mM ATPγS, 70 ug/mL BSA) followed by incubation for 3 hours in a 25 °C water bath. This pre-formed RecA* mixture was then mixed 1:1 with LexA in 1X control buffer (70 mM Tris-HCl, pH 7.5, 150 mM NaCl, 10 mM MgCl2, 2 mM TCEP, 70 ug/mL BSA) for the following final reactant concentrations: [RecA] = 1.00-1.25 µM, [GGT18] = 100-250 nM, and [LexA] = 2.5 µM. At each time point, 10 µL of each reaction was added to 10 µL 2X Laemmli buffer and boiled at 95 °C to stop the reaction. For the zero-minute time point, 5 µL of each RecA* and LexA mix were independently added to the 10 µL stop buffer. 10 µL of each sample was run on a 15% SDS-PAGE gel for analysis via Coomassie staining.

In cleavage assays testing the effects of SOS operator, final [LexA] was reduced to 1.5 µM and 20 µM of the respective operator was included in the LexA mixture and preincubated at 25 °C for 30 minutes prior to mixing with RecA*.

For assays requiring more sensitive quantification, the reaction mixture also contained 50 nM of a LexA variant containing a CF488A maleimide dye (Thermo) conjugated to an introduced Cys residue at position 201 (LexA-CF). These gels were then imaged on a Typhoon imager using the Cy2 laser line at a fixed 250 PMT voltage. Construct labeling was done as previously described and has been shown to have no impact on LexA autoproteolysis rates at this position^44^. Additionally, to maintain steady state assumptions, the concentration of RecA (final) was reduced to 400 nM and the concentration of GGT18 (final) was reduced to 30 nM.

### RecA Filamentation Assay

Activated RecA* was prepared in the same way as above, additionally containing 2 nM of a 3’-FAM labeled GGT18 oligo. Anisotropy was measured after the 3-hour activation on a Panvera Beacon2000 with a 490 nm excitation filter and a 525 nm emission filter (Farrand Controls). Blank solutions were prepared with no labeled GGT18 and control reactions for RecA WT were prepared by withholding ATPγS.

### LexA Binding Assay

To measure LexA binding, LexA Q161δ (LexA-δ) was spiked into the pre-formed RecA* to a final concentration of 50 nM. These samples were incubated for 30 minutes at room temperature and then anisotropy was measured on a Panvera Beacon2000 using a 387 ±11 nm excitation filter and a 448 ±20 nm emission filter (Edmond Optics). Blank solutions were prepared with a LexA storage buffer spike-in.

For binding competition experiments, unlabeled LexA K156A or QM were spiked into the pre-formed RecA* alongside the LexA Q161δ, spanning a final concentration range of unlabeled LexA of 140 nM to 18.3 µM. Data was fit using GraphPad Prism v 9.5.1 with the nonlinear least squares fit to the [Inhibitor] vs. response (three parameter) model, plotting the 95% confidence intervals of the fit (**Fig. 4B**).

### Statistical Analysis of Quantitative Biochemical Assays

For all quantitative analyses, the scale parameters (the normalized means for each data set) were compared using Bayesian analysis. Due to having only three replicates, we use a T-distribution joint posterior distribution rather than the Gaussian noise model, using a uniform prior:

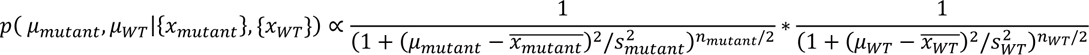

Where *μ_mutant_* and *μ_WT_* are the estimates of their true values, *S_mutant_* and *S_WT_* are the variances, 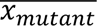 and 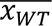 are the sample means, and *n_mutant_* and *n_WT_* are the number of replicates for the mutant and WT datasets respectively. Substituting the following:

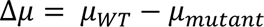

Gives:

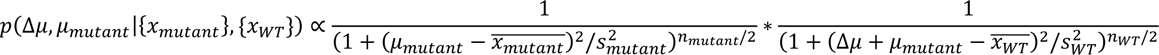

Normalizing the posterior probability density function (pdf) and plotting the corresponding cumulative density function (cdf) allows for the estimation of the likelihood that the value of Δ*μ* is greater than zero using a 95% cutoff (cdf of y = 0.05).

### UniProt Accession Codes for Proteins Referenced in this Study

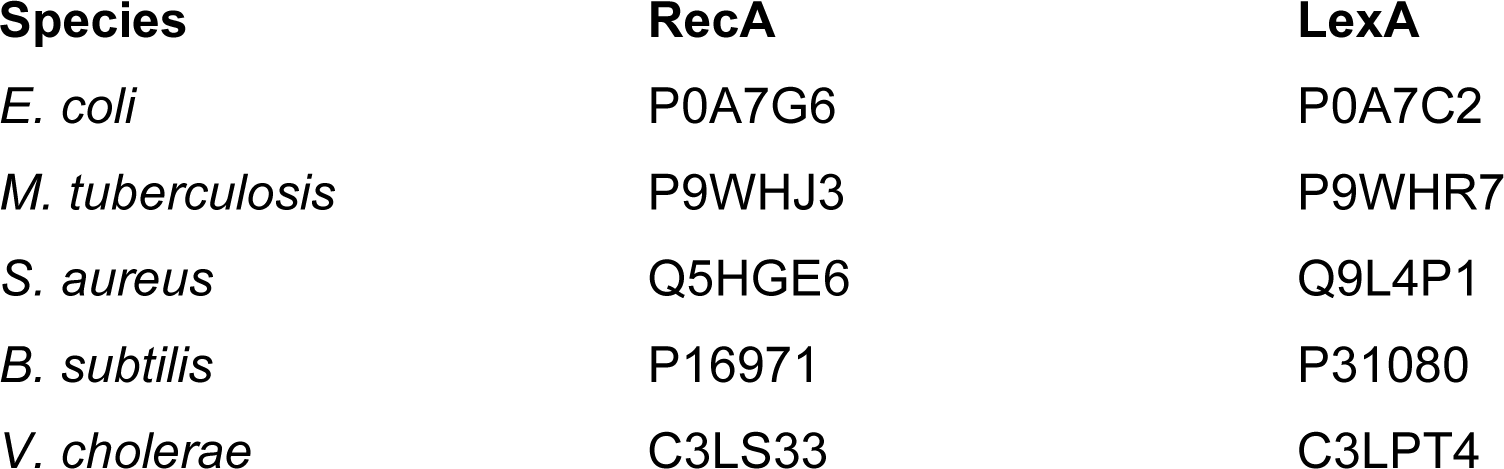
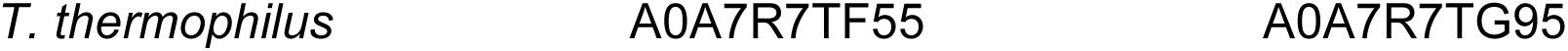

| Species | RecA | LexA |
| --- | --- | --- |
| *E. coli* | P0A7G6 | P0A7C2 |
| *M. tuberculosis* | P9WHJ3 | P9WHR7 |
| *S. aureus* | Q5HGE6 | Q9L4P1 |
| *B. subtilis* | P16971 | P31080 |
| *V. cholerae* | C3LS33 | C3LPT4 |

### Data Availability

All raw data and images, as well as plasmid sequences are available on request.

## Notes

### Competing Interest Statement

The authors have declared no competing interest.

## REFERENCES

1. Culyba, M.J., Mo, C.Y. & Kohli, R.M. Targets for Combating the Evolution of Acquired Antibiotic Resistance. Biochemistry 54, 3573–3582 (2015).

2. Blair, J.M., Webber, M.A., Baylay, A.J., Ogbolu, D.O. & Piddock, L.J. Molecular mechanisms of antibiotic resistance. Nat. Rev. Microbiol. 13, 42–51 (2015).

3. Darby, E.M. et al. Molecular mechanisms of antibiotic resistance revisited. Nat. Rev. Microbiol. 21, 280–295 (2023).

4. Maslowska, K.H., Makiela-Dzbenska, K. & Fijalkowska, I.J. The SOS system: A complex and tightly regulated response to DNA damage. Environ. Mol. Mutagen. 60, 368–384 (2019).

5. Memar, M.Y. et al. The central role of the SOS DNA repair system in antibiotics resistance: A new target for a new infectious treatment strategy. Life Sci. 262, 118562 (2020).

6. Podlesek, Z. & Žgur Bertok, D. The DNA Damage Inducible SOS Response Is a Key Player in the Generation of Bacterial Persister Cells and Population Wide Tolerance. Front. Microbiol. 11, 1785 (2020).

7. Recacha, E. et al. Quinolone Resistance Reversion by Targeting the SOS Response. MBio 8, 10.1128/mBio.00971-17 (2017).

8. Da Re, S. et al. The SOS response promotes qnrB quinolone-resistance determinant expression. EMBO Rep. 10, 929–933 (2009).

9. Mo, C.Y. et al. Systematically Altering Bacterial SOS Activity under Stress Reveals Therapeutic Strategies for Potentiating Antibiotics. mSphere 1, 163 (2016).

10. Revitt-Mills, S.A. et al. Defects in DNA double-strand break repair resensitize antibiotic-resistant Escherichia coli to multiple bactericidal antibiotics. Microbiology Open 11, e1316 (2022).

11. Ragheb, M.N. et al. Inhibiting the Evolution of Antibiotic Resistance. Mol. Cell 73, 157–165.e5 (2019).

12. Crane, J.K., Burke, S.R. & Alvarado, C.L. Inhibition of SOS Response by Nitric Oxide Donors in Escherichia coli Blocks Toxin Production and Hypermutation. Front. Cell. Infect. Microbiol. 11, 798136 (2021).

13. Wipperman, M.F. et al. Mycobacterial Mutagenesis and Drug Resistance Are Controlled by Phosphorylation-and Cardiolipin-Mediated Inhibition of the RecA Coprotease. Molecular Cell 72, 152–161.e7 (2018).

14. Brent, R. & Ptashne, M. Mechanism of action of the lexA gene product. Proc. Natl. Acad. Sci. U. S. A. 78, 4204–4208 (1981).

15. Fernandez De Henestrosa, A.R., et al. Identification of additional genes belonging to the LexA regulon in Escherichia coli. Mol. Microbiol. 35, 1560–1572 (2000).

16. Kamensek, S., Podlesek, Z., Gillor, O. & Zgur-Bertok, D. Genes regulated by the Escherichia coli SOS repressor LexA exhibit heterogeneous expression. BMC Microbiol. 10, 283–283 (2010).

17. Butala, M. et al. Interconversion between bound and free conformations of LexA orchestrates the bacterial SOS response. Nucleic Acids Res. 39, 6546–6557 (2011).

18. Bell, J.C. & Kowalczykowski, S.C. RecA: Regulation and Mechanism of a Molecular Search Engine. Trends Biochem. Sci. 41, 491–507 (2016).

19. Sankar, T.S., Wastuwidyaningtyas, B.D., Dong, Y., Lewis, S.A. & Wang, J.D. The nature of mutations induced by replication-transcription collisions. Nature 535, 178–181 (2016).

20. Paul, S., Million-Weaver, S., Chattopadhyay, S., Sokurenko, E. & Merrikh, H. Accelerated gene evolution through replication-transcription conflicts. Nature 495, 512–515 (2013).

21. Arana, M.E. & Kunkel, T.A. Mutator phenotypes due to DNA replication infidelity. Semin. Cancer Biol. 20, 304–311 (2010).

22. Ganai, R.A. & Johansson, E. DNA Replication-A Matter of Fidelity. Mol. Cell 62, 745–755 (2016).

23. Mo, C.Y., Birdwell, L.D. & Kohli, R.M. Specificity determinants for autoproteolysis of LexA, a key regulator of bacterial SOS mutagenesis. Biochemistry 53, 3158–3168 (2014).

24. Brent, R. Regulation and autoregulation by lexA protein. Biochimie 64, 565–569 (1982).

25. Slilaty, S.N., Rupley, J.A. & Little, J.W. Intramolecular cleavage of LexA and phage lambda repressors: dependence of kinetics on repressor concentration, pH, temperature, and solvent. Biochemistry 25, 6866–6875 (1986).

26. Giese, K.C., Michalowski, C.B. & Little, J.W. RecA-dependent cleavage of LexA dimers. J. Mol. Biol. 377, 148–161 (2008).

27. Kim, B. & Little, J.W. LexA and lambda Cl repressors as enzymes: specific cleavage in an intermolecular reaction. Cell 73, 1165–1173 (1993).

28. Rehrauer, W.M., Lavery, P.E., Palmer, E.L., Singh, R.N. & Kowalczykowski, S.C. Interaction of Escherichia coli RecA protein with LexA repressor. I. LexA repressor cleavage is competitive with binding of a secondary DNA molecule. J. Biol. Chem. 271, 23865–23873 (1996).

29. Luo, Y. et al. Crystal structure of LexA: a conformational switch for regulation of self-cleavage. Cell 106, 585–594 (2001).

30. Mohana-Borges, R. et al. LexA repressor forms stable dimers in solution. The role of specific DNA in tightening protein-protein interactions. J. Biol. Chem. 275, 4708–4712 (2000).

31. Neher, S.B., Flynn, J.M., Sauer, R.T. & Baker, T.A. Latent ClpX-recognition signals ensure LexA destruction after DNA damage. Genes Dev. 17, 1084–1089 (2003).

32. Friedman, N., Vardi, S., Ronen, M., Alon, U. & Stavans, J. Precise temporal modulation in the response of the SOS DNA repair network in individual bacteria. PLoS Biol. 3, e238 (2005).

33. Ni, M., Wang, S.Y., Li, J.K. & Ouyang, Q. Simulating the temporal modulation of inducible DNA damage response in Escherichia coli. Biophys. J. 93, 62–73 (2007).

34. Culyba, M.J., Kubiak, J.M., Mo, C.Y., Goulian, M. & Kohli, R.M. Non-equilibrium repressor binding kinetics link DNA damage dose to transcriptional timing within the SOS gene network. PLoS Genet. 14, e1007405 (2018).

35. Naiman, K., Philippin, G., Fuchs, R.P. & Pagès, V. Chronology in lesion tolerance gives priority to genetic variability. Proc. Natl. Acad. Sci. U. S. A. 111, 5526–5531 (2014).

36. Fujii, S. & Fuchs, R.P. A Comprehensive View of Translesion Synthesis in Escherichia coli. Microbiol. Mol. Biol. Rev. 84, 2 (2020).

37. Fuchs, R.P. Tolerance of lesions in E. coli: Chronological competition between Translesion Synthesis and Damage Avoidance. DNA Repair 44, 51–58 (2016).

38. Mo, C.Y. et al. Type III-A CRISPR immunity promotes mutagenesis of staphylococci. Nature 592, 611–615 (2021).

39. Marx, P. et al. Environmental stress perception activates structural remodeling of extant Streptococcus mutans biofilms. NPJ Biofilms Microbiomes 6, 17-z (2020).

40. Soares, A., Alexandre, K. & Etienne, M. Tolerance and Persistence of Pseudomonas aeruginosa in Biofilms Exposed to Antibiotics: Molecular Mechanisms, Antibiotic Strategies and Therapeutic Perspectives. Front. Microbiol. 11, 2057 (2020).

41. Adikesavan, A.K. et al. Separation of recombination and SOS response in Escherichia coli RecA suggests LexA interaction sites. PLoS Genet. 7, e1002244 (2011).

42. Kovacic, L. et al. Structural insight into LexA-RecA* interaction. Nucleic Acids Res. 41, 9901–9910 (2013).

43. Yu, X. & Egelman, E.H. The LexA repressor binds within the deep helical groove of the activated RecA filament. J. Mol. Biol. 231, 29–40 (1993).

44. Cory, M.B. et al. Engineered RecA Constructs Reveal the Minimal SOS Activation Complex. Biochemistry (2022).

45. Gao, B. et al. Structural basis for regulation of SOS response in bacteria. Proc. Natl. Acad. Sci. U. S. A. 120, e2217493120 (2023).

46. VanLoock, M.S. et al. Complexes of RecA with LexA and RecX differentiate between active and inactive RecA nucleoprotein filaments. J. Mol. Biol. 333, 345–354 (2003).

47. Chen, Z., Yang, H. & Pavletich, N.P. Mechanism of homologous recombination from the RecA-ssDNA/dsDNA structures. Nature 453, 489–484 (2008).

48. Egelman, E.H. & Stasiak, A. Structure of helical RecA-DNA complexes. Complexes formed in the presence of ATP-gamma-S or ATP. J. Mol. Biol. 191, 677–697 (1986).

49. Hostetler, Z.M., Cory, M.B., Jones, C.M., Petersson, E.J. & Kohli, R.M. The Kinetic and Molecular Basis for the Interaction of LexA and Activated RecA Revealed by a Fluorescent Amino Acid Probe. ACS Chem. Biol., 1127–1133 (2020).

50. Jaramillo, A.V.C., Cory, M.B., Li, A., Kohli, R.M. & Wuest, W.M. Exploration of inhibitors of the bacterial LexA repressor-protease. Bioorg. Med. Chem. Lett. 65, 128702 (2022).

51. Maso, L. et al. Nanobodies targeting LexA autocleavage disclose a novel suppression strategy of SOS-response pathway. Structure 30, 1479–1493.e9 (2022).

52. Mo, C.Y. et al. Inhibitors of LexA Autoproteolysis and the Bacterial SOS Response Discovered by an Academic-Industry Partnership. ACS Infect. Dis. 4, 349–359 (2018).

53. Cory, M.B., Hostetler, Z.M. & Kohli, R.M. Kinetic dissection of macromolecular complex formation with minimally perturbing fluorescent probes. Methods Enzymol. 664, 151–171 (2022).

54. Hörtnagel, K. et al. Saturation mutagenesis of the E. coli RecA loop L2 homologous DNA pairing region reveals residues essential for recombination and recombinational repair. J. Mol. Biol. 286, 1097–1106 (1999).

55. Courcelle, J., Khodursky, A., Peter, B., Brown, P.O. & Hanawalt, P.C. Comparative gene expression profiles following UV exposure in wild-type and SOS-deficient Escherichia coli. Genetics 158, 41–64 (2001).

56. Hostetler, Z.M. et al. Systematic Evaluation of Soluble Protein Expression Using a Fluorescent Unnatural Amino Acid Reveals No Reliable Predictors of Tolerability. ACS Chem. Biol. 13, 2855–2861 (2018).

57. Punjani, A., Rubinstein, J.L., Fleet, D.J. & Brubaker, M.A. cryoSPARC: algorithms for rapid unsupervised cryo-EM structure determination. Nat. Methods 14, 290–296 (2017).

58. Sanchez-Garcia, R. et al. DeepEMhancer: a deep learning solution for cryo-EM volume post-processing*. Commun*. Biol. 4, 874–1 (2021).

59. Naydenova, K. & Russo, C.J. Measuring the effects of particle orientation to improve the efficiency of electron cryomicroscopy. Nat. Commun. 8, 629–3 (2017).

60. Zivanov, J. et al. New tools for automated high-resolution cryo-EM structure determination in RELION-3. Elife 7, 10.7554/eLife.42166 (2018).

61. Jumper, J. et al. Highly accurate protein structure prediction with AlphaFold. Nature 596, 583–589 (2021).

62. Pettersen, E.F. et al. UCSF ChimeraX: Structure visualization for researchers, educators, and developers. Protein Sci. 30, 70–82 (2021).

63. Emsley, P., Lohkamp, B., Scott, W.G. & Cowtan, K. Features and development of Coot. Acta Crystallogr. D Biol. Crystallogr. 66, 486–501 (2010).

64. Moriarty, N.W., Grosse-Kunstleve, R.W. & Adams, P.D. electronic Ligand Builder and Optimization Workbench (eLBOW): a tool for ligand coordinate and restraint generation. Acta Crystallogr. D Biol. Crystallogr. 65, 1074–1080 (2009).

65. Liebschner, D. et al. Macromolecular structure determination using X-rays, neutrons and electrons: recent developments in Phenix. Acta Crystallogr. D. Struct. Biol. 75, 861–877 (2019).

66. Afonine, P.V. et al. Real-space refinement in PHENIX for cryo-EM and crystallography. Acta Crystallogr. D. Struct. Biol. 74, 531–544 (2018).

67. Croll, T.I. ISOLDE: a physically realistic environment for model building into low-resolution electron-density maps. Acta Crystallogr. D. Struct. Biol. 74, 519–530 (2018).

68. Williams, C.J. et al. MolProbity: More and better reference data for improved all-atom structure validation. Protein Sci. 27, 293–315 (2018).

69. Afonine, P.V. et al. New tools for the analysis and validation of cryo-EM maps and atomic models. Acta Crystallogr. D. Struct. Biol. 74, 814–840 (2018).

