## Supplemental Information for "The LexA-RecA* structure reveals a lock-and-key mechanism for SOS activation"

### SUPPLEMENTARY INFORMATION

**Figure S1:** Cryo-EM analysis pipeline………………………………………………. 1-2
**Figure S2:** Characteristics of the EM density and model…………………………. 3-4
**Figure S3:** Alkaline autoproteolysis rates of each tested LexA variant………… 5-6
**Figure S4:** Discrimination between operator-bound LexA and free LexA by…… RecA* 7
**Figure S5:** Analysis of potential charge-charge interactions between RecA*….. and LexA 8-9
**Figure S6:** Allosteric binding pocket on LexA and species variation……………. 9
**Figure S7:** Biochemical analysis of RecA3x mutant filamentation and LexA….. binding 10
**Figure S8:** SDS-PAGE of LexA constructs………………………………………… 10

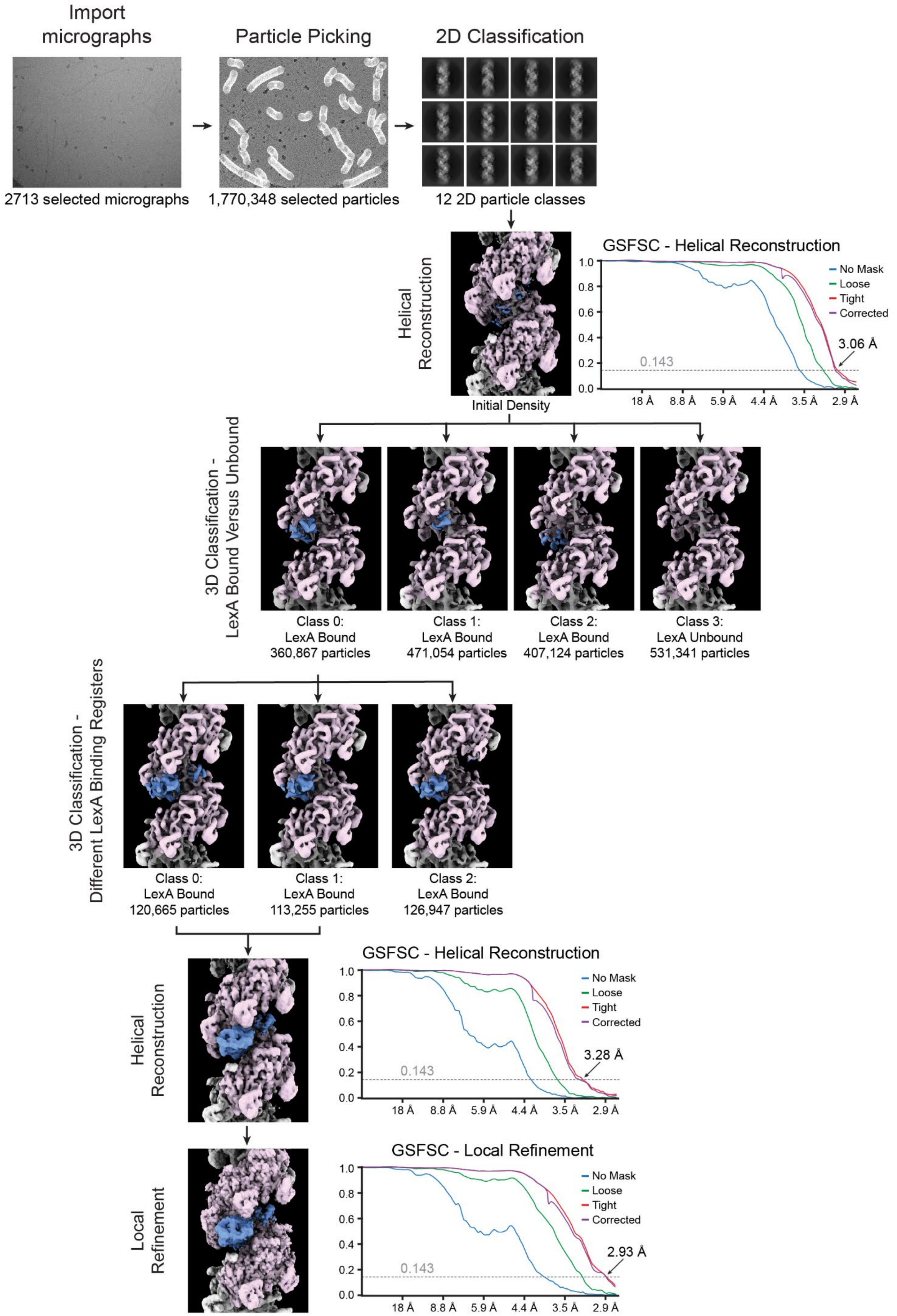
Import micrographs
Particle Picking
2D Classification
2713 selected micrographs
1,770,348 selected particles
12 2D particle classes
Helical Reconstruction
Initial Density
GSFSC - Helical Reconstruction
No Mask
Loose
Tight
Corrected
3.06 Å
0.143
1.0
0.8
0.6
0.4
0.2
0.0
18 Å
8.8 Å
5.9 Å
4.4 Å
3.5 Å
2.9 Å
3D Classification - LexA Bound Versus Unbound
Class 0: LexA Bound 360,867 particles
Class 1: LexA Bound 471,054 particles
Class 2: LexA Bound 407,124 particles
Class 3: LexA Unbound 531,341 particles
3D Classification - Different LexA Binding Registers
Class 0: LexA Bound 120,665 particles
Class 1: LexA Bound 113,255 particles
Class 2: LexA Bound 126,947 particles
Helical Reconstruction
GSFSC - Helical Reconstruction
No Mask
Loose
Tight
Corrected
3.28 Å
0.143
Local Refinement
GSFSC - Local Refinement
No Mask
Loose
Tight
Corrected
2.93 Å
0.143

**Figure S1. Cryo-EM analysis pipeline.** The flow of data from the collected and filtered micrographs through the final local refinement is shown. Each labeled step includes relevant information for the partitioning of data at each junction. For each refinement and reconstruction step, the FSC curve generated by CryoSPARC is shown.

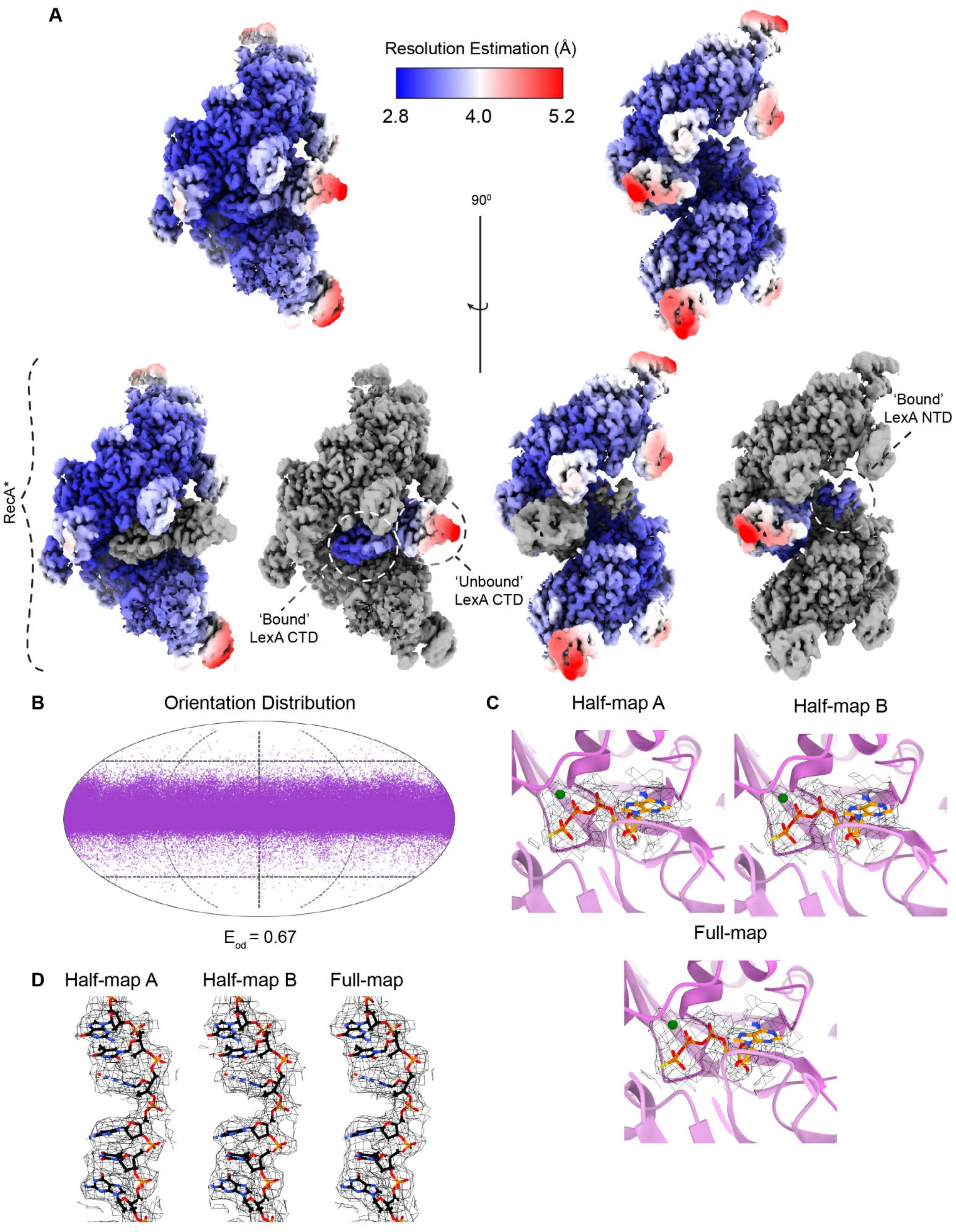
A
Resolution Estimation (Å)
2.8
4.0
5.2
90⁰
RecA*
'Bound'
LexA CTD
'Unbound'
LexA CTD
'Bound'
LexA NTD
B
Orientation Distribution
$E_{od}$ = 0.67
C
Half-map A
Half-map B
Full-map
D
Half-map A
Half-map B
Full-map

**Figure S2. Characteristics of the EM density and model. A)** Final sharpened map colored by the estimated local resolution using Relion. The entire complex is shown at top, with the relevant sub-complex components at the bottom with RecA* and LexA labeled and colored relative to grayed out other components. **B)** Orientation distribution of the final particle stack as determined by cryoEF. Orientation efficiency, $E_{od}$ is given below. **C)** Closeup of the ATP binding pocket at the interface of two RecA protomers within the filament, showing the coordinated $Mg^{2+}$ ion in green. Density from the two independent half maps and the corresponding full map at a contour level of 0.203 and 0.172 respectively shown in mesh. **D)** Closeup of the bound ssDNA within the filament. Density from the two independent half maps and the corresponding full map at a contour level of 0.203 and 0.172 respectively shown in mesh.

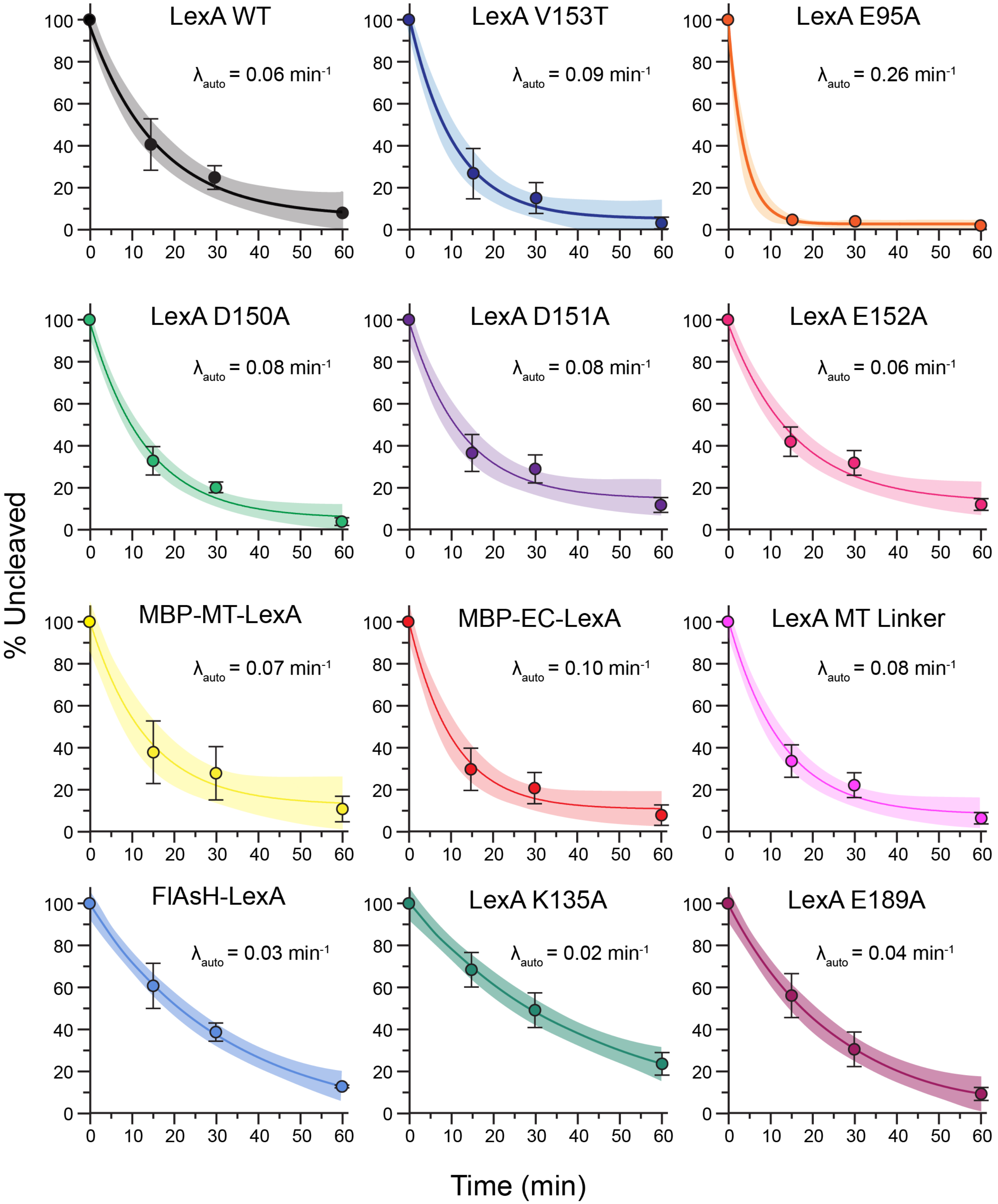


**Figure S3. Alkaline autoproteolysis rates of each tested LexA variant.** Data represents the mean from three replicates, with error bars denoting standard deviation. Data were fit to a single exponential decay (solid line) with 95% confidence intervals shown (shaded region). The best-fit value for the decay rate is shown on each graph.

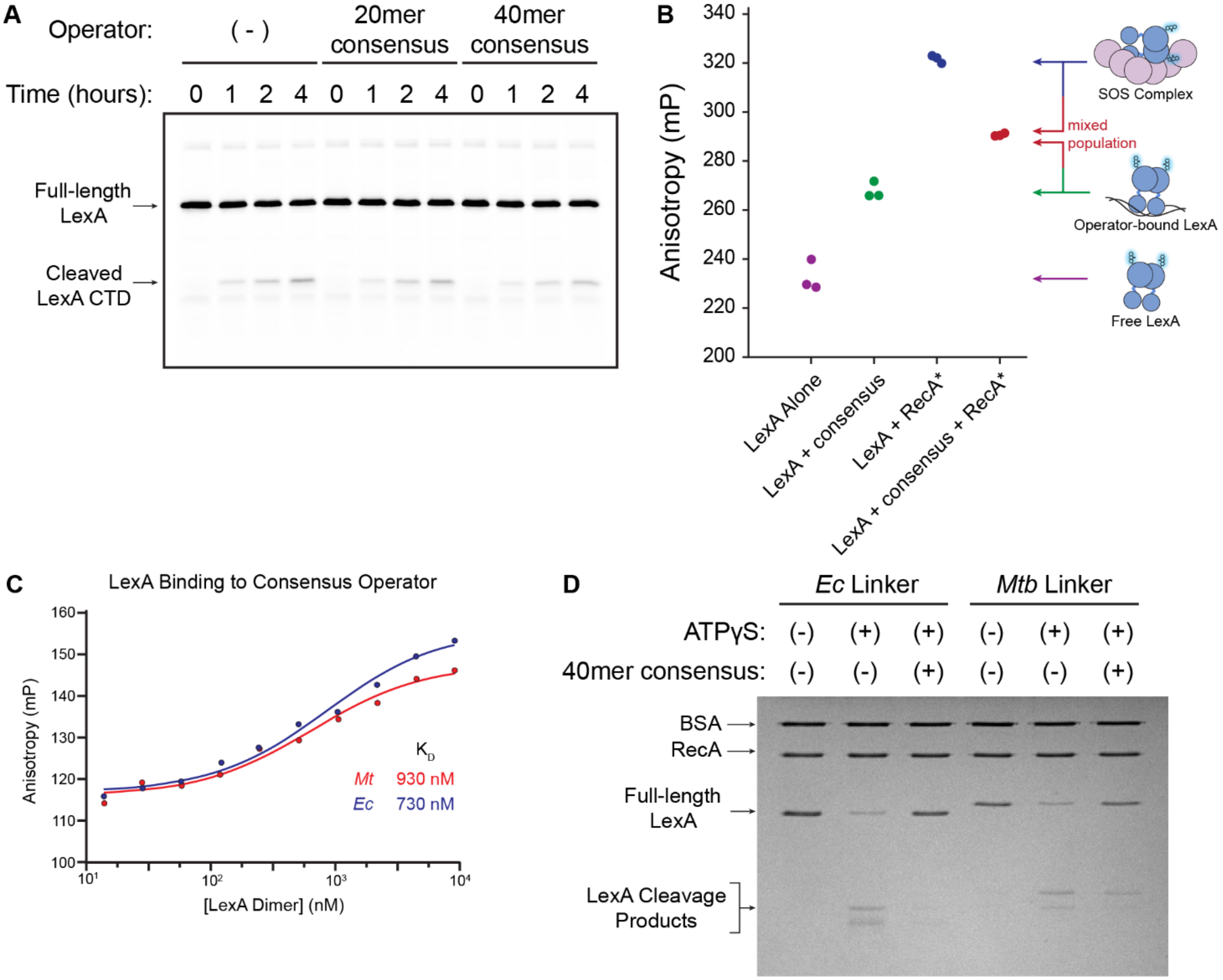


**Figure S4. Discrimination between operator-bound LexA and free LexA by RecA*.** **A)** SDS-PAGE gel of autoproteolysis of fluorescent LexA-CF variant at pH 7.5 in the absence of operator or in the presence of either 20 bp or 40 bp consensus operator. **B)** Fluorescence anisotropy of LexA-δ with various *in vitro* binding partners. Each data point represents a single replicate. The various contributing species to the observed anisotropic signal are given to the right. **C)** Equilibrium endpoint anisotropy titration of either *Ec* or *Mtb* LexA with FAM-labeled 40mer consensus operator. Data shows a single replicate and the solid line is a fit to a quadratic equation, using a fixed [operator] of 1 nM. **D)** SDS-PAGE analysis of RecA*-dependent cleavage of *E. coli* LexA with either an *Ec* or *Mtb* inter-domain linker when incubated with consensus operator.

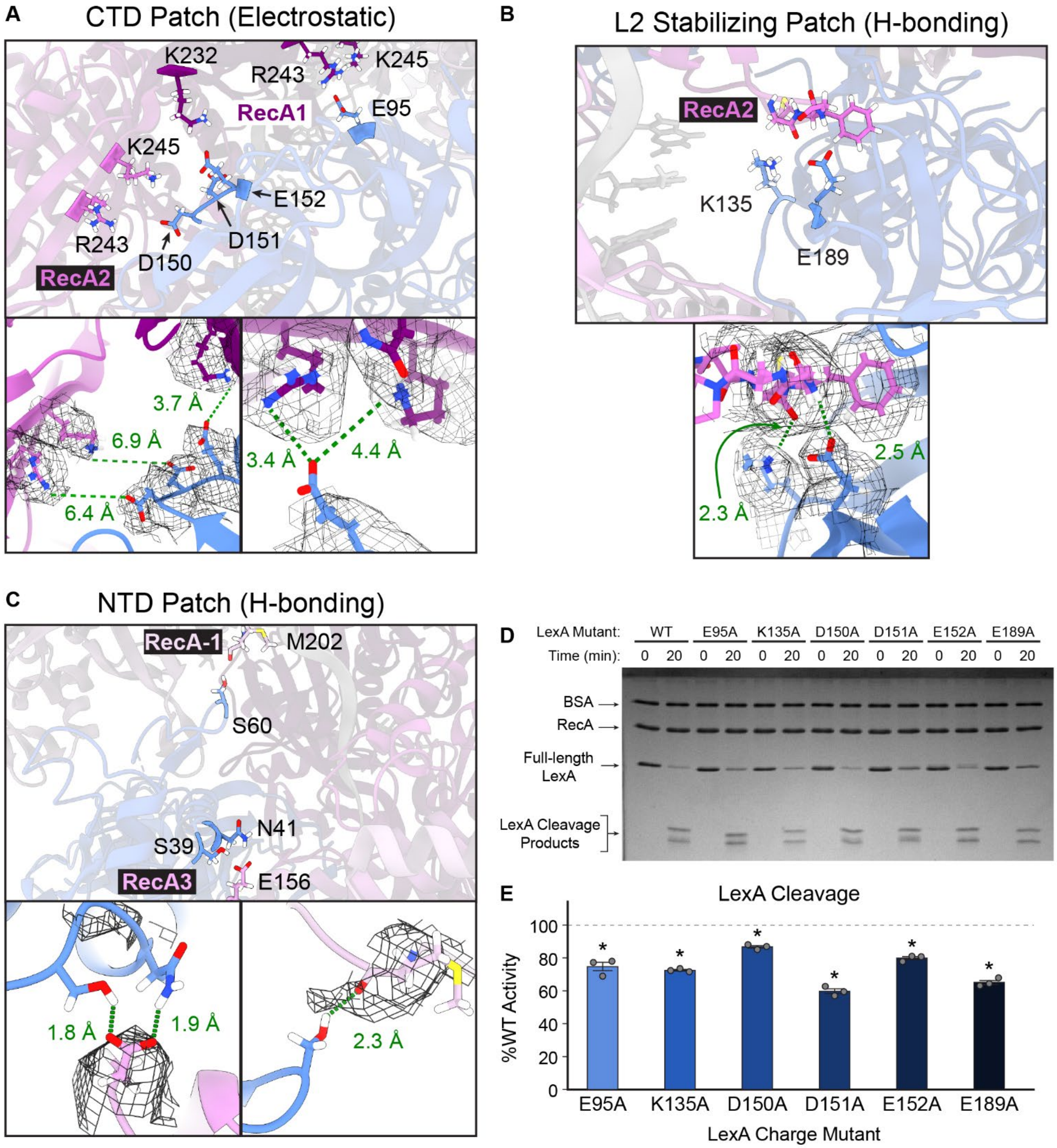


**Figure S5. Analysis of potential charge-charge interactions between RecA* and LexA. A-C)** Three different sections within the interaction interface that provide potential charge-charge interactions. Each of the RecA protomers within the three consecutive RecA units providing a majority of the contacts are highlighted and labeled in shades of pink to purple. The bound LexA monomer is shown in blue. Insets highlight the distances between interacting residues in green. **D)** Representative SDS-PAGE analysis of RecA*-dependent cleavage of each LexA charge mutant. **E)** Quantified LexA cleavage expressed as a percentage of WT LexA rate (normalized to 100% shown by the dotted line). Each bar represents the mean from three replicates and error bars denote standard deviation. * denotes mutants with a >95% posterior likelihood of being less than WT.

**A**

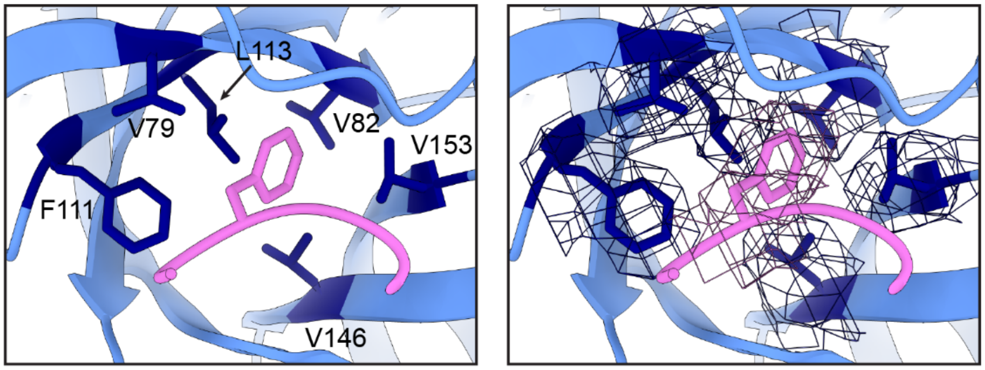


**B**

| | LexA L2 Pocket | | | | | | | RecA L2 Loop |
|---|---|---|---|---|---|---|---|---|
| Position: | 79 | 82 | 111 | 113 | 146 | 153 | | 198..................208 |
| | V | V | F | L | V | V | *E. coli* | K I G V M F G N P E T |
| | L | I | F | L | A | A | *M. tuberculosis* | K I G V M F G S P E T |
| | I | V | F | L | V | A | *S. aureus* | K V G V M F G N P E T |
| | I | V | F | L | V | A | *B. subtilis* | K V G V M F G N P E T |
| | I | V | F | L | V | V | *V. cholerae* | K I G V M F G N P E T |
| | V | V | F | L | A | T | *T. thermophilus* | K V G V T Y G N P E T |

**Figure S6. Allosteric binding pocket on LexA and species variation. A)** Surface and cartoon representations of the SOS complex model, as shown in Fig. 2. RecA F203 (pink) is bound to LexA (blue) within the hydrophobic pocket formed by the highlighted LexA residues (purple). The map density is shown on the right as a mesh surface. **B)** Sequence alignment of LexA and RecA proteins from select different species, showing the LexA hydrophobic pocket residues (left) and a subsection of the RecA L2 loop (right). F203 is highlighted in pink.

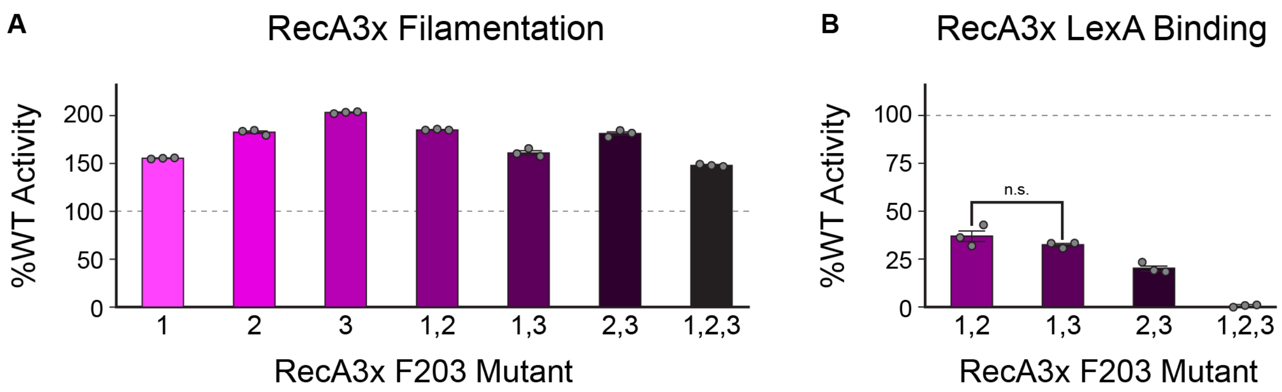


**Figure S7. Biochemical analysis of RecA3x mutant filamentation and LexA binding. A)** RecA3x mutant filamentation expressed as a percentage of RecA3x WT anisotropy (normalized to 100% shown by the dotted line) in the FAM-ssDNA binding assay. Data show the means of three replicates, with error bars denoting standard deviation. **B)** RecA3x mutant binding to LexA-δ expressed as a percentage of RecA3x WT anisotropy (normalized to 100% shown by the dotted line) in the LexA binding assay. Data show the means of three replicates, with error bars denoting standard deviation. Unless otherwise noted, all mutants had a >95% posterior likelihood of being less than both RecA3x WT and each other for LexA binding percentage.

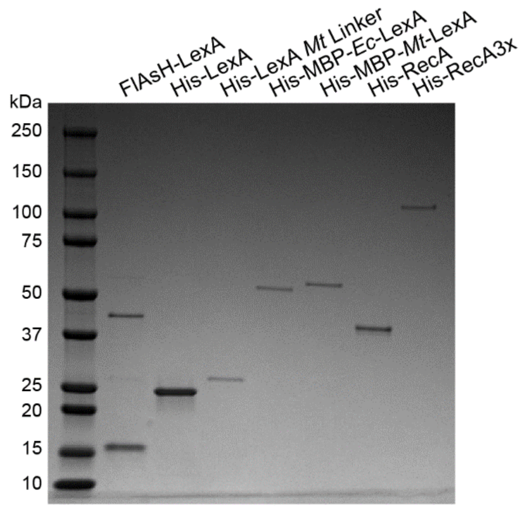


**Figure S8. LexA constructs.** Shown is the SDS-PAGE gel including various LexA constructs evaluated in this study with the molecular weights of the ladder labeled at left.
